# Defects in tissue-resident macrophages lead to smaller eardrums and abnormal neurovascular networks with increased middle-ear infection

**DOI:** 10.64898/2026.01.13.699360

**Authors:** Yunpei Zhang, Pingting Wang, Lingling Neng, George Burwood, Kushal Sharma, Allan Kachelmeier, Xiaorui Shi

## Abstract

The tympanic membrane (TM), or eardrum, is essential for hearing. Macrophages, the primary innate immune cells, are densely distributed in the eardrum after birth, especially near blood vessels and nerve fibers. During postnatal development, neonatal tissue-resident macrophages (TRMs) gradually polarize, and their population declines. These dynamic changes closely parallel the maturation of neurovascular networks. What are the precursors of the early ‘wave’ of TRMs? Are they critical for postnatal TM development? Using fate mapping, single-cell RNA sequencing, and macrophage depletion, this study reveals for the first time that postnatal eardrum TRMs are heterogeneous, originating mainly from embryonic myeloid lineages, with increasing input from postnatal monocyte progenitor-derived cells. Single-cell RNA sequencing identifies gene signatures vital for vascular and neuronal development. Depleting TRMs results in smaller eardrums, disrupted vascular-neuronal networks, and increased risk of middle-ear infection. This study offers new insight into how the innate immune system supports TM maturation and protects against middle-ear infection during a critical postnatal window.

**Teaser:** Tissue-resident macrophages guide eardrum neurovascular maturation and help prevent middle-ear abnormalities.

## Introduction

Tissue-resident macrophages (TRMs) are essential components of the innate immune system, exhibiting diverse phenotypes and functional specialization across nearly all tissues and organs, including the tympanic membrane (TM), or eardrum (*1, 2*). Recent studies have shown that TRMs are predominantly established during embryogenesis, with considerable heterogeneity in their origin, maintenance, and function depending on the tissue microenvironment (*3, 4*). Fate-mapping and barcoding studies indicate that many TRMs are established embryonically from yolk sac-derived erythro-myeloid progenitors (EMPs) that differentiate through pre-macrophage intermediates, with variable degrees of postnatal monocyte input depending on tissue context (*5–8*). In addition, subsets of tissue macrophages can be progressively supplemented or replaced after birth by myeloid lineages derived from granulocyte-monocyte progenitors (GMPs) and bone marrow hematopoiesis, particularly in tissues undergoing remodeling or exposed to environmental challenge (*7, 8*). In general, these macrophages contribute broadly to tissue development, homeostasis, and inflammation (*9–11*). However, how TRMs develop and function within the eardrum remains poorly understood, despite the eardrum’s constant exposure to environmental challenge and its essential role in hearing.

The eardrum is a thin, semi-transparent oval membrane separating the external auditory canal from the middle ear. It plays a critical role in hearing by vibrating in response to sound waves and transmitting mechanical energy to the auditory ossicles, which then stimulate sensory hair cells that convert these signals into neural impulses. A combination of lever gain through the middle ear, and the surface area ratio of the eardrum to the footplate of the terminal ossicle, the stapes, facilitates mechanical impedance matching between the air in the auditory canal and the fluid of the cochlea, vastly increasing the efficiency of sound transmission to the inner ear (*12*). Structurally, the eardrum consists of two distinct regions: the pars tensa, the main vibratory component (*13*), and the pars flaccida. It is densely vascularized and innervated, features that are essential for its metabolic demands and sensory function (*14*). However, despite the known importance of TRMs in other tissues and the recognized complexity of the eardrum, the developmental trajectory and functional roles of TRMs within the eardrum remain unexplored. This gap limits our understanding of how immune cells contribute to normal eardrum maturation and how defects in these processes may increase susceptibility to middle-ear abnormalities.

In this study, we employed three established transgenic mouse models to trace the developmental origin of eardrum TRMs. Our findings indicate that eardrum TRMs are largely established by embryonic myeloid lineages (yolk sac + fetal liver), with progressive acquisition of postnatal input from the GMP lineage during maturation. To define TRM diversity and potential functions, we performed single-cell RNA sequencing (scRNA-seq) on macrophages isolated from postnatal day 15 (P15) eardrums and identified gene signatures associated with vascular and neural development. To assess their functional relevance, we utilized a macrophage ablation model in neonatal mice, which demonstrated that embryonic TRMs are critical for proper vascular remodeling, peripheral nerve maturation, and overall eardrum growth. Notably, TRM depletion led to increased incidence of middle ear abnormalities (fluid and fibrosis) and smaller eardrums with abnormal vascular and neuronal architecture. We used *in vivo* OCT structural imaging to detect frequent middle-ear fluid accumulation and fibrotic changes in macrophage-depleted ears, supporting a role for TRMs in maintaining middle-ear homeostasis during development. While structural abnormalities of the eardrum have long been reported in clinical settings (*15, 16*), the cellular and developmental mechanisms underlying these defects remain elusive. Our findings provide new insights into how TRMs contribute to the structural and functional maturation of the eardrum, uncovering a previously unrecognized role for innate immune cells in auditory physiology. In this work, the scientific question is whether TRMs coordinate eardrum neurovascular maturation while limiting middle-ear abnormalities during a vulnerable postnatal window.

## Results

### Eardrum macrophage dynamics from neonatal to young adult stages

*Cx3cr1^EGFP^* reporter mouse model is frequently used across various organs to study macrophages (*17*), as *Cx3cr1* encodes a chemokine receptor that is specifically expressed in mononuclear phagocytes, serving as a distinct marker for identifying macrophages (*18*). In this study, we discovered the abundance of CX3CR1^+^ cells resident in the eardrums of *Cx3cr1^EGFP^*mouse. In neonatal mice, the majority of CX3CR1^+^ cells in eardrum are unpolarized, exhibit a branched morphology, and are primarily located in the pars flaccida, the manubrial portion of the pars tensa, and the annulus. As the animals mature, the CX3CR1^+^ cells become noticeably elongated, polarized, and orient radially toward the center of the manubrial portion (Fig. 1A-B). These features are more clearly visualized under high magnification (Fig. 1A, *lower panel* and Fig. 1B). The density of CX3CR1^+^ cells decreases as the animals reach adulthood (Fig. 1C). To confirm that the CX3CR1^+^ cells are macrophages, we employed an additional macrophage marker, F4/80 (*19*), to double-label the CX3CR1^+^ cells, as shown in Fig. 1D. All CX3CR1^+^ macrophages are F4/80^+^.

**Fig. 1.**
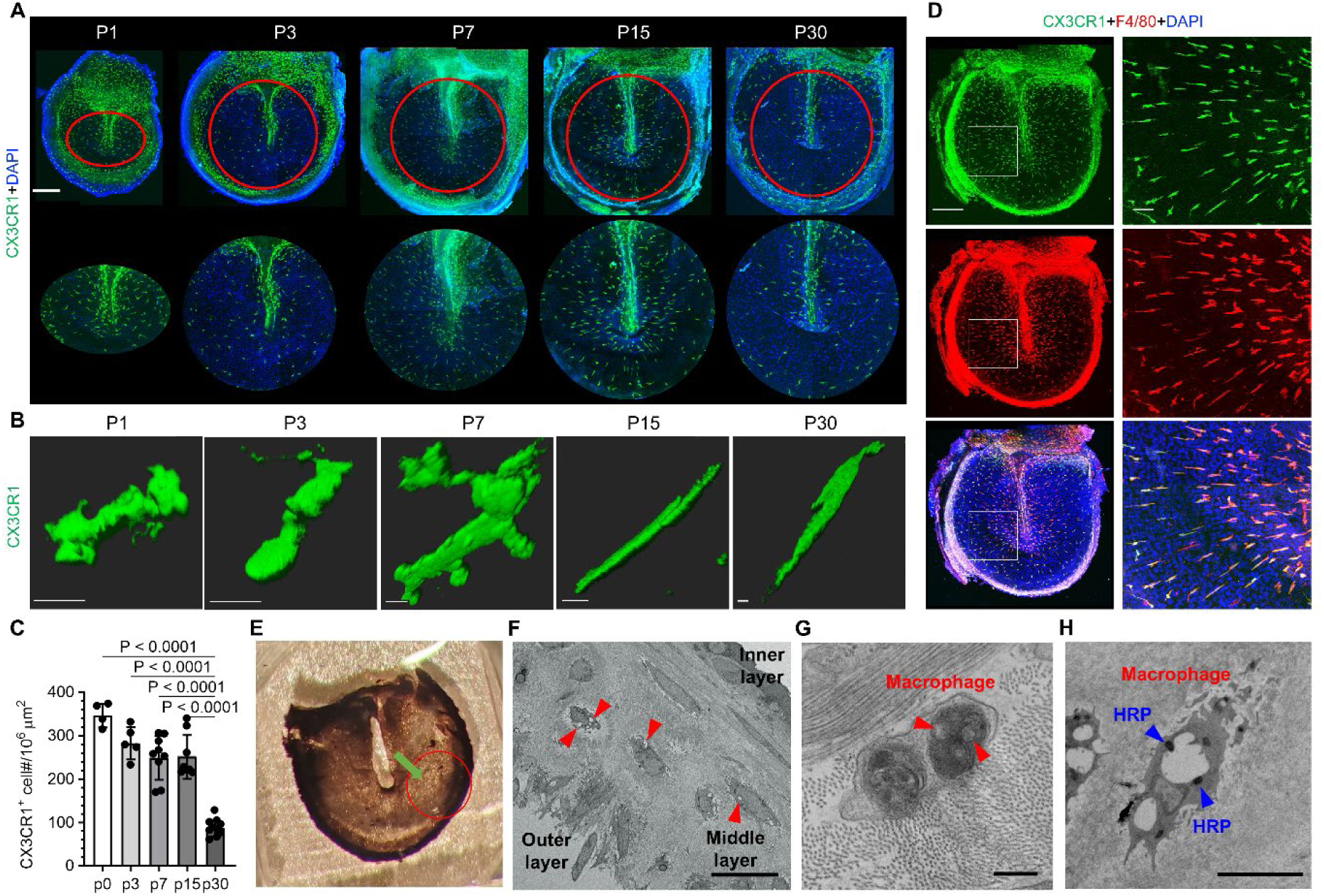
Dynamics of TRMs in *CX3CR1^EGFP^* mice from newborn to young adults. (**A**) Representative confocal projection images show the distribution pattern of CX3CR1+ cells in the eardrum at various postnatal stages: postnatal day 1 (P1), day 3 (P3), day 7 (P7), day 15 (P15), and day 30 (P30). The images show the pars tensa regions at low magnification *(upper panel)*, higher magnification *(lower panel)*. Scale bar: 300 μm. (**B**) High magnification images of the morphology and orientation of macrophages during eardrum development from birth to young adult. Scale bar: 5 μm. (**C**) The quantification of CX3CR1+ cells density across developmental stages: P1 (n=4), P3 (n=5), P7 (n=9), P15 (n=8), and P30 (n=10). Data are presented as mean ± SEM. Statistical significance was assessed using one-way ANOVA with Tukey’s test (F (4, 31) = 43.25). (**D**) Representative confocal images demonstrate co-localization of EGFP-tagged CX3CR1 expression with the macrophage marker F4/80. Scale bar: 300 μm. (**E-H**) show that macrophages situate in the middle layer of tympanic membrane. Specifically, **E** and **F** show TEM images obtained under low magnification, showing the location of eardrum examined and three layers of eardrum structure and macrophage distribution identified by in the middle fibrous layer of the tympanic membrane. TRMs are identified by pinocytotic vesicles **G** and uptake HRP pointed by red arrowheads under high magnification **H**. Scale bars: 15 μm (**F**), 200 nm (**G**), 4 μm (**H**).

TM is composed of three distinct layers: an outer epidermal layer, a middle fibrous layer, and an inner mucosal layer of epithelium (*20*). At the ultrastructural level, we observed that macrophages are in the middle layer of the eardrum. This layer contains densely organized collagen fibers, subdivided into an external radial layer and an internal circular layer, along with fibroblasts, nerve fibers, and capillaries (*21*). The macrophages were morphologically identified by their highly developed surface projections, including their long, slender filopodia containing lysosomes, phagosomes, and macro-pinosomes, and by their abundance of polyribosomes(*22, 23*). We have also used a conventional method to facilitate identification of macrophages by administering horseradish peroxidase (HRP) type II to the animals as macrophages can uptake the HRP due to their phagocytotic activity(*24*). Figs. 1E–H show macrophages taking up HRP type II 18 hours prior to animal sacrifice under transmission electron microscopy (TEM). Taken together, our data demonstrate that eardrum macrophages change in response to local environmental changes after birth.

### Eardrum-resident macrophages are heterogeneous and primarily originate from embryonic sources

To determine the developmental origin of eardrum TRMs after birth and their dynamics during postnatal development, we employed two inducible Cre-lox fate-mapping models (*Csf1r-Mer-iCre-Mer;R26^tdTomato^* and *Cx3cr1^CreER^;R26^tdTomato^*) and a constitutive monocyte-lineage tracing model (*Ms4a3^Cre^;R26^tdTomato^*) (Fig. 2A). The population dynamics from these fate-mapping approaches are shown in Fig. 2B-D and quantified in Fig. 2F-I. The overall combined trend is summarized in Fig. 2J. *Csf1r*- and *Cx3cr1*-based embryonic pulses were used to mark embryonic myeloid lineages that give rise to tissue-resident macrophages, whereas *Ms4a3^Cre^*was used to quantify macrophages from the granulocyte-monocyte progenitor (GMP) lineage during postnatal life.

**Fig. 2.**
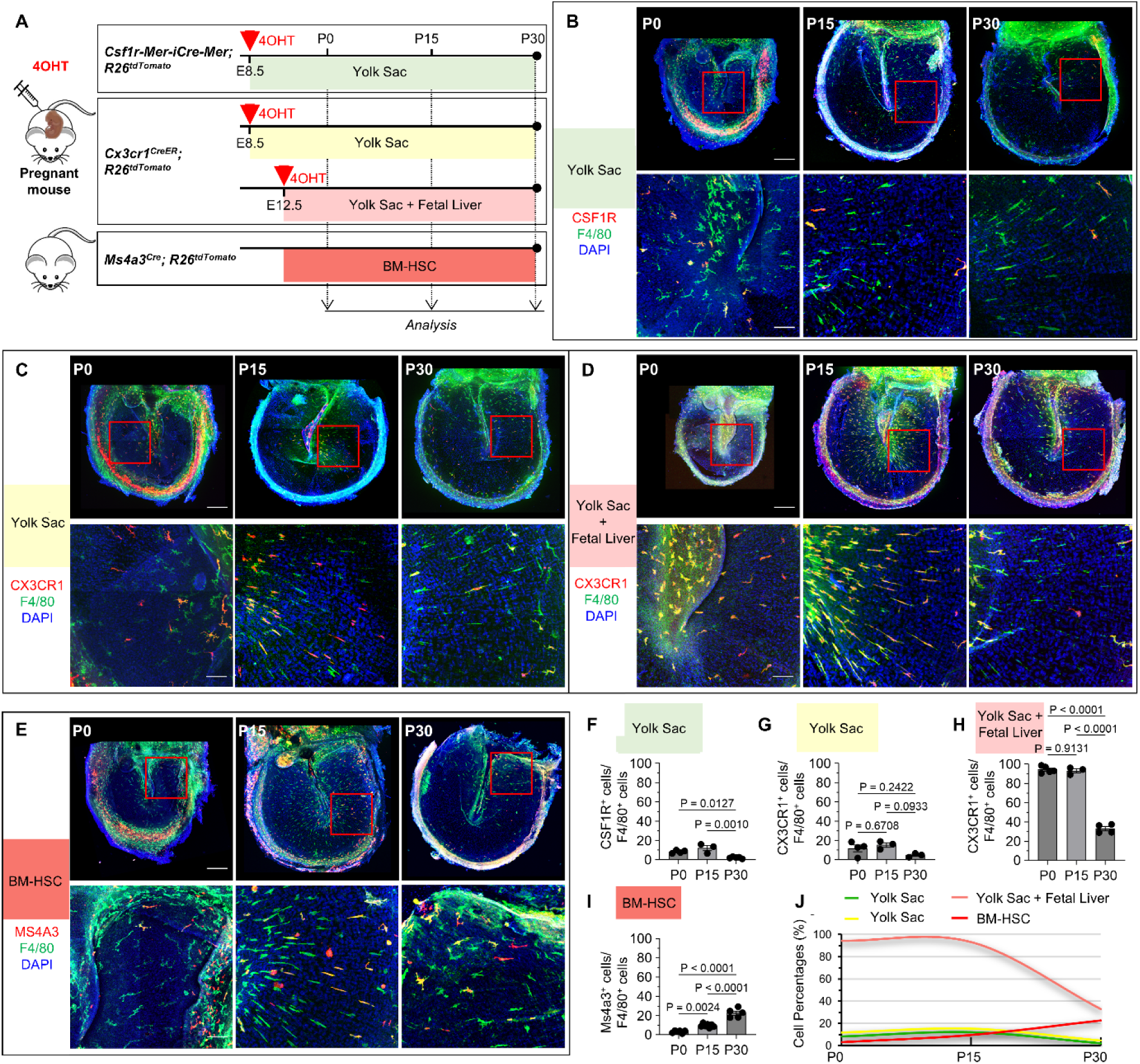
Postnatal eardrum macrophages are heterogeneous and originate from multiple sources. (**A**) Schematic representation of eardrum macrophage lineage tracing using various Cre animal models. (**B-E**) Representative confocal images demonstrating eardrum TRMs originating from YS (**B, C**), YS + fetal liver (**D**) and BM-HSC derived monocytes (**E**) mice at P0, P15, and P30. Scale bars: 300 μm (*upper panels*), 50 μm (*lower panels*). (**F-I**) Population dynamics of TRM derived from YS-CSF1R (**F**, P1 (n=4), P15 (n=3), P30 (n=5)), YS-CX3CR1(**G**, P1 (n=4), P15 (n=3), P30 (n=3)), YS + Fetal liver (**H**, P1 (n=5), P15 (n=3), P30 (n=4)), and BM-derived monocytes (**I**, P1 (n=6), P15 (n=7), P30 (n=5)). (**J**) Summary of overall lineage-tracing trends across panels F–I. Data were presented as mean ± SEM, statistical significance was determined by One-way ANOVA with Tukey’s test (F_YS-CSF1R_ (2, 9) = 16.27, F_YS-CX3CR1_ (2, 7) = 3.246, F_YS + Fetal liver_ (2, 9) = 325.4, F_BM-HSC_ (2, 15) = 66.40).

Specifically, to trace yolk sac (YS)-derived embryonic myeloid lineages that populate tissues and differentiate into TRMs, we administered 4-hydroxytamoxifen (4OHT) at embryonic day 8.5 (E8.5) in these two inducible Cre models, a timepoint corresponding to YS-derived myeloid cells that enter tissues and mature through a pre-macrophage stage. Our results indicated only a sparse presence of *Csf1r^tdTomato^* and *Cx3cr1^tdTomato^* macrophages in postnatal eardrums (Fig. 2B, C, F, and G), indicating that an E8.5 pulse labels only a subset of the embryonically established TRM pool in this tissue.

To determine embryonic CX3CR1⁺ myeloid cells at mid-gestation, we administered 4OHT to *Cx3cr1^CreER^; R26^tdTomato^*mice at E12.5, an embryonic state when the fetal liver becomes the primary hematopoietic organ. CX3CR1 is expressed by multiple embryonic myeloid populations at E12.5, this pulse labels YS-derived macrophage lineage cells that have already entered tissues and fetal liver-derived myeloid cells, so these labeling captures mixed embryonic contributions. Notably, we observed a substantial increase in labeled macrophages in the eardrum (Fig. 2D, H), supporting a major contribution of embryonically established macrophage lineages to the postnatal eardrum TRM pool. However, these E12.5-labeled macrophage populations declined in number by P30 (Fig. 2H), consistent with a developmental shift in TRM maintenance and/or increasing replacement by postnatal myeloid inputs during maturation.

In addition, we assessed postnatal contributions from the GMP lineage using *Ms4a3^Cre^; R26^tdTomato^* mice. *Ms4a3* is primarily expressed in early myeloid lineage cells, specifically GMPs, and *Ms4a3* fate mapping labels granulocytes and monocytes(*25*). At birth (P0), only a small number of *Ms4a3*-lineage macrophages were present in the pars tensa of eardrum, whereas a larger population was observed in adjacent bone tissues (annulus), likely within osteoblasts and osteocytes (Fig. 2E). Notably, the number of *Ms4a3*-lineage macrophages increased steadily during the postnatal period from P1 to P30 (Fig. 2I). Collectively, these findings suggest that TRMs are heterogeneous and originate from multiple sources.

### Postnatal remodeling of eardrum vasculature and innervation

Alongside TRMs dynamics, we noticed postnatal vascular remodeling within the eardrum. In neonatal mice, the vascular network forms a plexus-like structure, resembling early anastomotic vessels, referred to here as ‘nascent’ blood vessels. These immature vessels are densely concentrated in specific areas, including the pars flaccida, the umbo of the malleus, and the annulus (Fig. 3A), areas that also exhibit high densities of CX3CR1^+^ macrophages (Fig. S1). The nascent vascular network undergoes significant morphological refinement as development progresses, becoming less branched and narrower in diameter. These developmental transitions are more clearly visualized at higher magnification (Fig. 3B). Quantitative analysis confirmed a progressive reduction in overall vascular density (Fig. 3C) and a concomitant decrease in vessel diameter (Fig. 3D) from early postnatal stages to adulthood. The vascular regression pattern is similarly and consistently observed in *NG2DsRedBAC* transgenic mice, which expresses an optimized red fluorescent protein variant (DsRed.T1) controlled by the mouse NG2 (*Cspg4*) promoter/enhancer, a mark for vascular pericytes (Fig. S2).

**Fig. 3.**
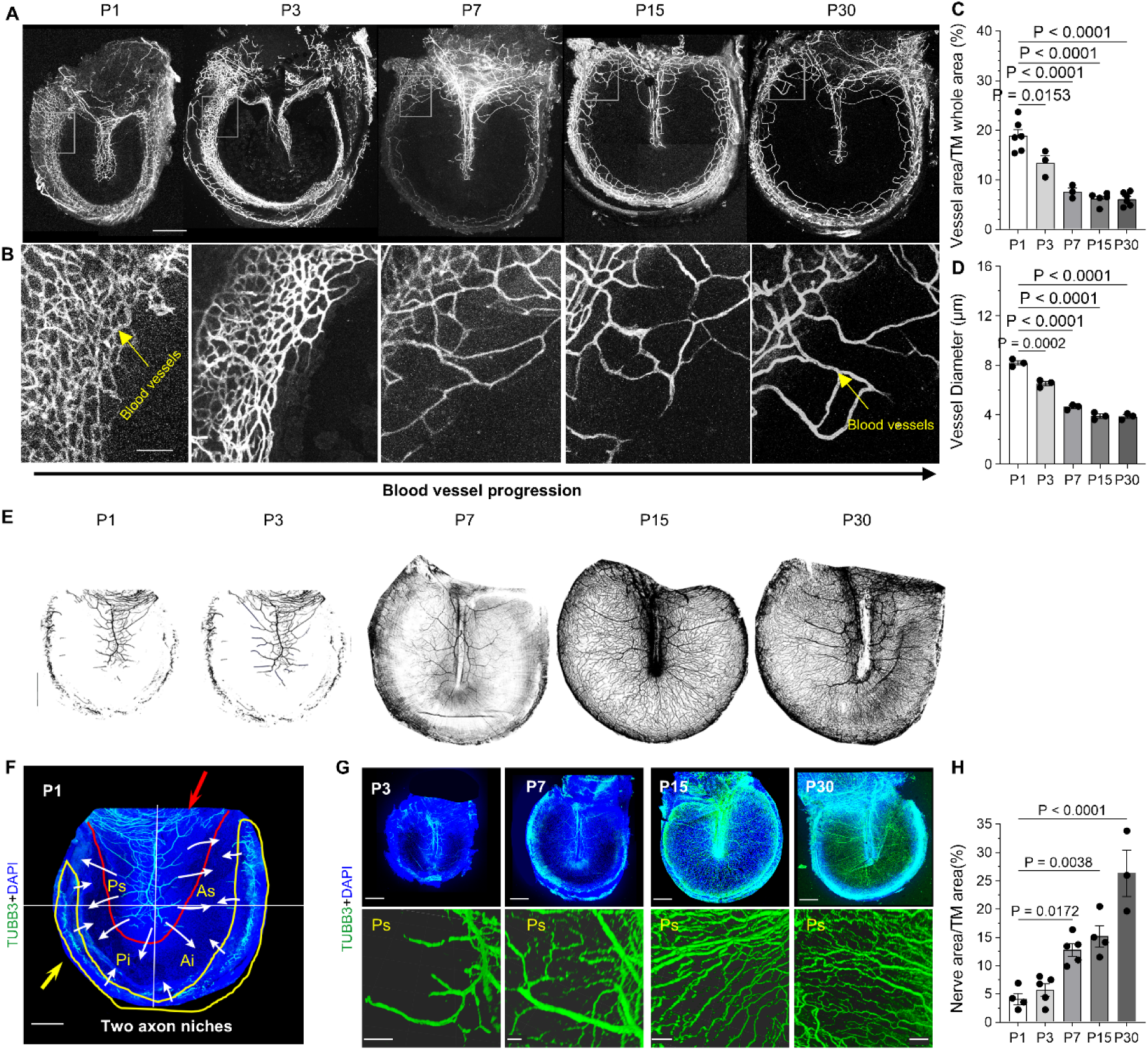
Postnatal development of blood vessels and peripheral nerve fiber fibers in the eardrum. (**A, B**) Confocal images demonstrate vascular network development in the eardrum of postnatal *NG2^DsRed^* mice. Scale bars: 300 μm (**A**),75 μm (**B**). (**C**) Quantitative analysis of vascular density at P1 (n=6), P3 (n=3), P7 (n=3), P15 (n=5), and P30 (n=6). Data were presented as mean ± SEM, statistical significance was assessed by One-way ANOVA with Tukey’s test (F (4, 18) = 36.85). (**D**) Quantitative analysis of vessel diameters at P1, P3, P7, P15, and P30 (3 replicates for each group, 10 vessels were randomly selected per sample within a defined region of interest). Data were presented as mean ± SEM, statistical significance was assessed through One-way ANOVA with Tukey test (F (4, 10) = 141.2). (**E**) Illustration of never fiber development. (**F**) Representative images illustrating nerve fiber development labeled by βIII-tubulin (TUBB3) in the eardrum. Scale bars: 150 μm (**E**), 300 μm (**F** *upper*), 50 μm (**F** *lower*). (**G**) Statistical quantification of nerve fiber density across developmental stages: P1 (n=4), P3 (n=5), P7 (n=5), P15 (n=4), P30 (n=3). Data were presented as mean ± SEM, statistical significance was calculated through One-way ANOVA with Tukey’s test (F (4, 16) = 21.62).

In parallel, we examined the postnatal development of peripheral nerve fibers in the eardrum as illustrated in (Fig. 3E). At birth, two distinct axon “niches” are evident: one in the manubrial region of the pars tensa and another near the annulus (Fig. 3F). At P3, the peripheral axons are still immature, and neurite outgrowth in the pars tensa remains incomplete. By P7, nerve fibers—particularly those originating from the manubrial region—begin to extend across the membrane. At P15, axons from both regions integrate, covering most of the eardrum, although with slightly lower density. By P30, peripheral nerves are fully developed, exhibiting dense, well-organized networks (Fig. 3G). Quantitative analysis confirmed a significant increase in nerve fiber density following birth (Fig. 3H). Together, the increased density of nerve fibers and the reduction of blood vessels suggest an adaptive response of the vascular and nervous systems during the early postnatal period, which may be crucial for eardrum function.

### Macrophages regulate postnatal eardrum neurovascular development

The next question we asked: what potential morphological and molecular connections exist between macrophages and the neurovascular system?

To examine the special relationship between TRMs, blood vessels, and nerve fibers during postnatal development of the eardrum, we performed high-resolution imaging analyses. Notably, TRMs exhibited a spatial distribution closely aligned with both developing nerve fibers and blood vessels in the eardrum (Fig. 4A). Using dual fluorescent reporter mice, in which TRMs are labeled with EGFP, while vascular cells, such as pericytes (essential for vascular development, stability and maturation(*26*)), are labeled with DsRed (see materials and methods), we observed dynamic interactions between TRMs and pericytes during the postnatal developmental period (Fig. 4B). Additionally, TRMs were found adjacent to elongating nerve fibers in the eardrum (Fig. 4C). Their location may indicate a potential role in coordinating neurovascular development. For instance, macrophages at P1 were observed to position ahead of axonal fronts, suggesting a potential guidance role. By P7, several macrophages elongate or branch along the sprouting axons, forming a three-dimensional scaffold. At P15, the orientation of macrophages becomes polarized toward the axons, and in some cases, macrophages are found directly attached to the nerve fibers.

**Fig. 4.**
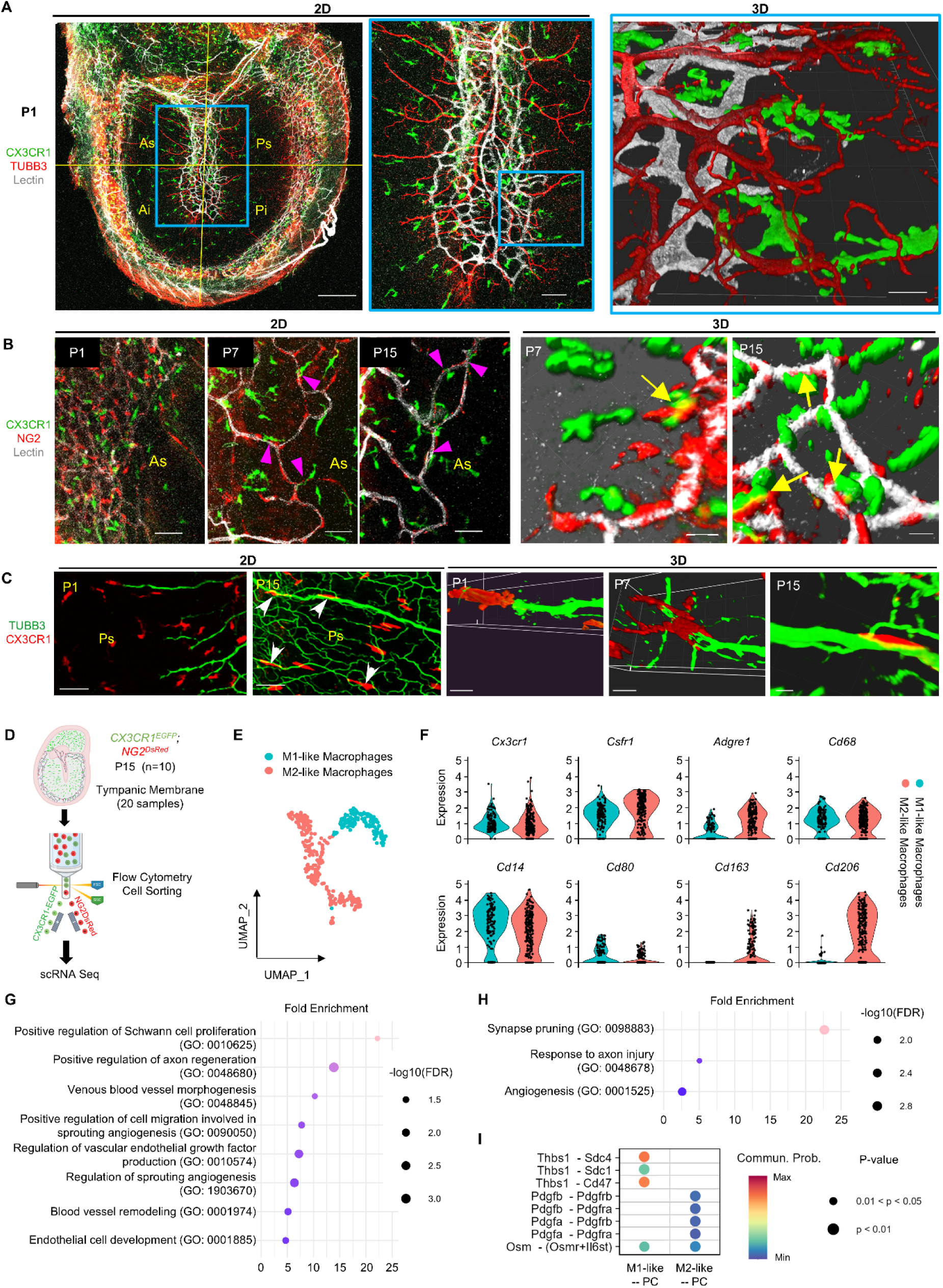
Spatial relationship between TRMs and the neurovascular network, highlighting macrophage-enriched gene clusters associated with bioprocesses in neuronal and vascular development. (**A**) Representative 2D and 3D confocal images showing a close association between eardrum TRMs, nerve fibers, and blood vessels at P1. Scale bars: 200 μm (*left*), 50 μm (*middle*), 20 μm (*right*). (**B**) Representative 2D (*left*) and 3D (*right*) images of macrophage interactions with blood vessel and pericytes (arrows) during postnatal development. Scale bars: 50 μm. (**C**) Representative 2D (*left*) and 3D (*right*) images of macrophage interactions with nerve fiber during postnatal development. Arrows, macrophages align with nerve fibers. Scale bars: 50 μm (*left*), 10 μm (*right*). (**D, E**) scRNA-seq analysis identifying distinct CX3CR1^+^ cell clusters from *CX3CR1^EGFP^; NG2^DsRed^* mouse eardrums (20 eardrums from 10 animals). (**F**) Violin plot of canonical macrophage markers (*Cx3Cr1, Csfr1, Adgre1, Cd68*) and subtype markers distinguishing M1-like and M2-like macrophages. (**G, H**) GO Biological Process enrichment (top 500 genes) in M1-like and M2-like macrophages related to neurovascular development. (**I**) Bubble plot (CellChat) of predicted ligand-receptor interactions that related to angiogenesis from macrophages to PCs; Dot size indicates p-value; color reflects communication probability.

To further investigate the molecular relationship between TRMs and the vascular-neural system during postnatal development, we conducted scRNA-seq on TRMs and pericytes isolated from the eardrums of P15 *CX3CR1^EGFP^; NG2^DsRed^* mice. Transcriptomic profiling identified distinct CX3CR1⁺ cell clusters (Fig. 4D-E). Quality-control metrics and filtering criteria are shown in Fig. S3. Canonical macrophage markers (*Cx3cr1*, *Csf1r*, F4/80/*Adgre1*, and *Cd68*) confirmed macrophage identity across clusters, while subtype markers separated the CX3CR1⁺ macrophages into M1-like and M2-like populations (Fig. 4F). To infer potential functions of these macrophage subsets, we performed GO Biological Process enrichment on the top 500 enriched genes in each subtype. M1-like macrophages were strongly associated with programs supporting neurovascular growth and remodeling (Fig. 4G), including positive regulation of Schwann cell proliferation (GO:0010625) and positive regulation of axon regeneration (GO:0048680), along with multiple vascular-development terms such as venous blood vessel morphogenesis (GO:0048845), positive regulation of cell migration involved in sprouting angiogenesis (GO:0090050), regulation of vascular endothelial growth factor production (GO:0010574), regulation of sprouting angiogenesis (GO:1903670), blood vessel remodeling (GO:0001974), and endothelial cell development (GO:0001885). In contrast, M2-like macrophages were enriched for pathways consistent with tissue maintenance and neural refinement following injury (Fig. 4H), including synapse pruning (GO:0098883) and response to axon injury (GO:0048678), while also retaining a vascular component through enrichment of angiogenesis (GO:0001525). Together, these GO profiles suggest that M1-like macrophages are biased toward pro-remodeling, pro-sprouting neurovascular functions, whereas M2-like macrophages emphasize neural circuit refinement and injury response alongside angiogenic support. Finally, CellChat analysis predicted macrophage-to-pericyte communication through ligand-receptor pairs associated with angiogenesis, notably involving Oncostatin M (Osm), a cytokine previously implicated in vascular remodeling (*27*) (Fig. 4I). Collectively, these data support a regulatory role for CX3CR1⁺ macrophage subsets in coordinating both angiogenic and neurogenic programs during postnatal eardrum development.

### Macrophage depletion impairs eardrum growth, neurovascular maturation, and increases susceptibility to middle ear infection

To investigate the role of postnatal macrophages in eardrum development, we utilized classical macrophage ‘suicide’ approach to avoid diphtheria toxin (DT) as its fusion toxins potential caused toxicity including ototoxicity, myocarditis and neuropathy in neonatal mice (*28–30*). Accordingly, we employed a liposome-mediated macrophage depletion strategy. Neonatal mice received injections of clodronate liposomes or control PBS liposomes via the facial vein. Within 24 hours, eardrum-resident macrophages effectively took up the liposomes, and by 72 hours, they metabolized them (Fig. S4A). To ensure consistent depletion, we administered additional intraperitoneal (*i.p*.) injections every three days from P2 until P15. This protocol resulted in nearly complete depletion of eardrum macrophages by P15. The sustained depletion of eardrum macrophages throughout the treatment window was confirmed (Fig. 5A and B). Compared to the control groups, mice that underwent macrophage depletion showed increased mortality and reduced body weight by P28 (Fig. 5C and D). Consistent with impaired maturation, macrophage-depleted mice exhibited smaller eardrums with reduced nerve fiber and vascular network development (Fig. 5E, H). Quantitative analysis confirmed these phenotypes, revealing significant differences in macrophage density, eardrum size, vessel branching, and nerve fiber density (Fig. 5B and I, Fig. S5B and C). In addition to the reduced size and branching, the blood vessels in the macrophage-depleted mice displayed abnormal accumulation of pericytes and signs of ongoing vascular sprouting, suggesting a failure to achieve full vascular maturation (Fig. 5J). These findings indicate that TRMs are crucial for coordinating the structural and functional maturation of both the vascular and neural components of the developing eardrum.

**Fig. 5.**
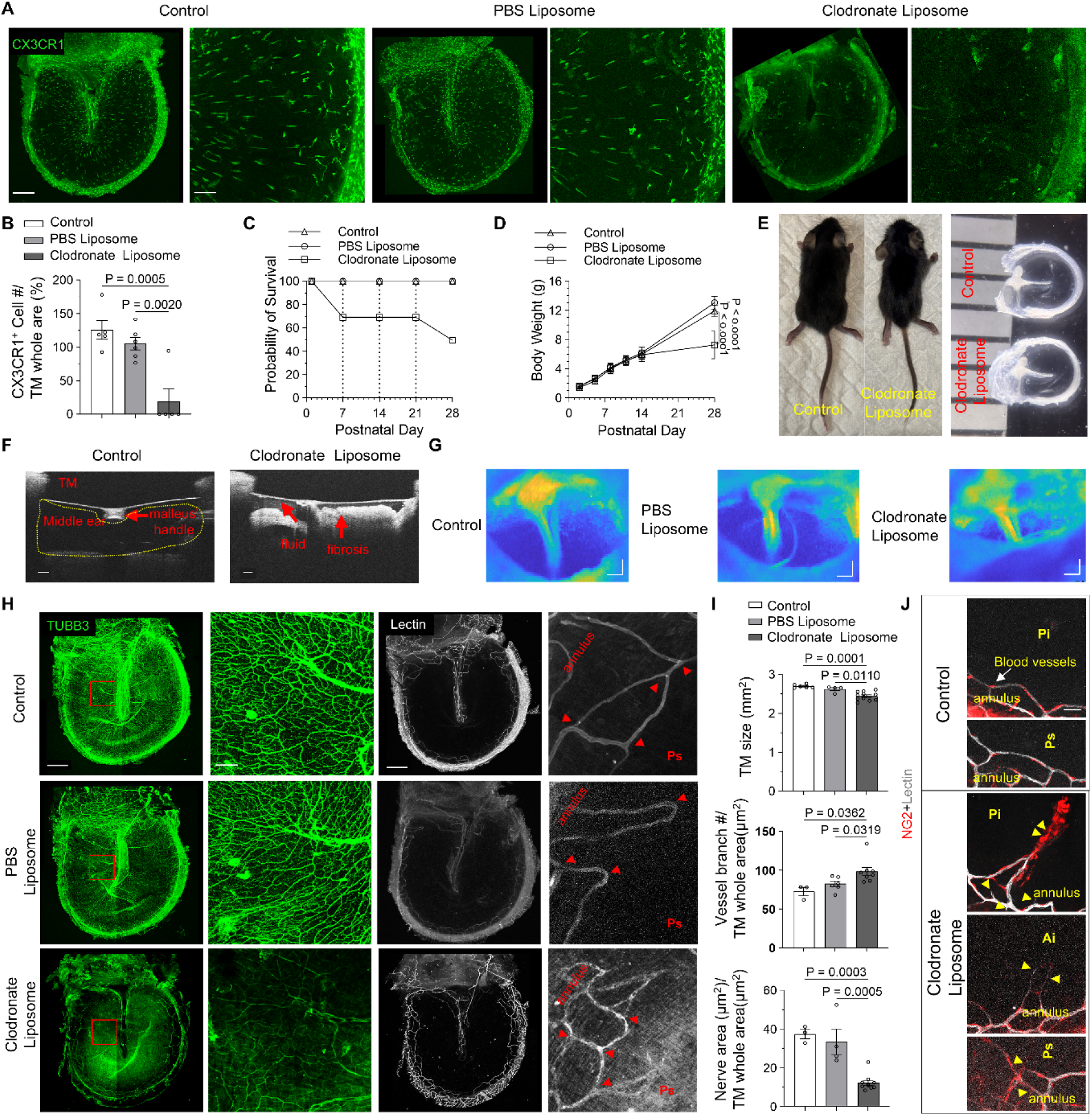
Macrophage depletion impairs growth and eardrum development. (**A**) Macrophage distribution at P15 in normal, liposome control, and macrophage-depleted animals. Scale bars: 300 μm (*low magnification*), 50 μm (*high magnification*). (**B**) Quantitative analysis of macrophage density at P15 (control n=5, PBS liposome n=6, Clodronate liposome n=5). Data were presented as mean ± SEM, statistical significance was calculated through One-way ANOVA with Tukey’s test (F (2, 13) = 19.26). (**C**) Kaplan-Meier survival curves comparing normal control, liposome control, and macrophage-depleted animals (control n=9, PBS liposome n=9, Clodronate liposome n=17). (**D**) General animal body weight analysis among different groups (control n=5, PBS liposome n=7, Clodronate liposome n=11). Data were presented as mean ± SEM, statistical significance was calculated through Two-way ANOVA with Šidak’s test (F_Interaction_ (10, 120) = 22.85, F_Raw Factor_ (5, 120) = 364.4, F_Column Factor_ (2, 120) = 24.55). (**E**) Representative images showing general animal size (*left*) and eardrum morphology (*right*) of normal and macrophage-depleted animals at P15. Scale bar (*right panel*): 1 mm. (**F**) *In vivo* 2D OCT B-scans of the eardrum reveal a notable accumulation of fluid in the middle ear and signs of fibrosis in the macrophage-depleted group (*lower panel*). Scale bar: 150 μm. (**G**) En face OCT images demonstrate field of view of the eardrum *in vivo.* Scale bar: 150 μm. (**H**) Distribution of nerve fibers and blood vessels in different groups. Scale bars: 300 μm (*low magnification*), 50 μm (*high magnification*). Arrowheads indicate vessel branching. (I) Quantitative analysis of eardrum size (control n=6, PBS liposome n=4, Clodronate liposome n=11), vessel branching density (control n=3, PBS liposome n=6, Clodronate liposome n=7), and nerve fiber density (control n=3, PBS liposome n=4, Clodronate liposome n=10) at P15. Data were presented as mean ± SEM, statistical significance was calculated through One-way ANOVA with Tukey’s test (F_TM size_ (2, 18) = 16.23, F_vessel branch density_ (2, 9) = 11.47, F_nerve fiber density_ (2, 14) = 21.76). **j** Confocal images depicting vessel sprouting and pericyte accumulation in macrophage-depleted versus control mice at P15. Ai: anteroinferior quadrant; Pi: posteroinferior quadrant; Ps: posterior superior quadrant. Scale bar: 25 μm.

To further confirm the impact of TRMs on the neurovascular network while minimizing systemic effects, local liposome injections were performed in neonatal mice. For local depletion, clodronate liposomes were administered via subcutaneous injection (*s.c.*) into the right pinna at a dose of 5 μL per gram of body weight (Fig. 6A). The initial dose was given to P2 mice, with subsequent injections every three days until P15 to sustain macrophage depletion during early postnatal development. Compared to systemic depletion, locally treated mice showed normal survival (data not shown) and did not show overt deficits in overall body growth (Fig. 6B and C). However, the injected pinna (auricle) fails to stand upright (Fig. 6B, *left panel*, red arrow, Fig. S6A) and is smaller in size (Fig. 6B, *upper right panels*). TM appeared mildly reduced in size and showed visible surface debris at the injection site (Fig. 6B, *lower right panels*, red arrow; 6E, *left panels*; 6F, *upper left panel*). Consistently reduced nerve fiber and blood vessel development, and abnormal accumulation of pericytes are also observed (Fig. 6E–G). Together, these data indicate that local TRM depletion is sufficient to impair neurovascular maturation and is associated with middle-ear abnormalities, while largely sparing systemic growth.

**Fig. 6.**
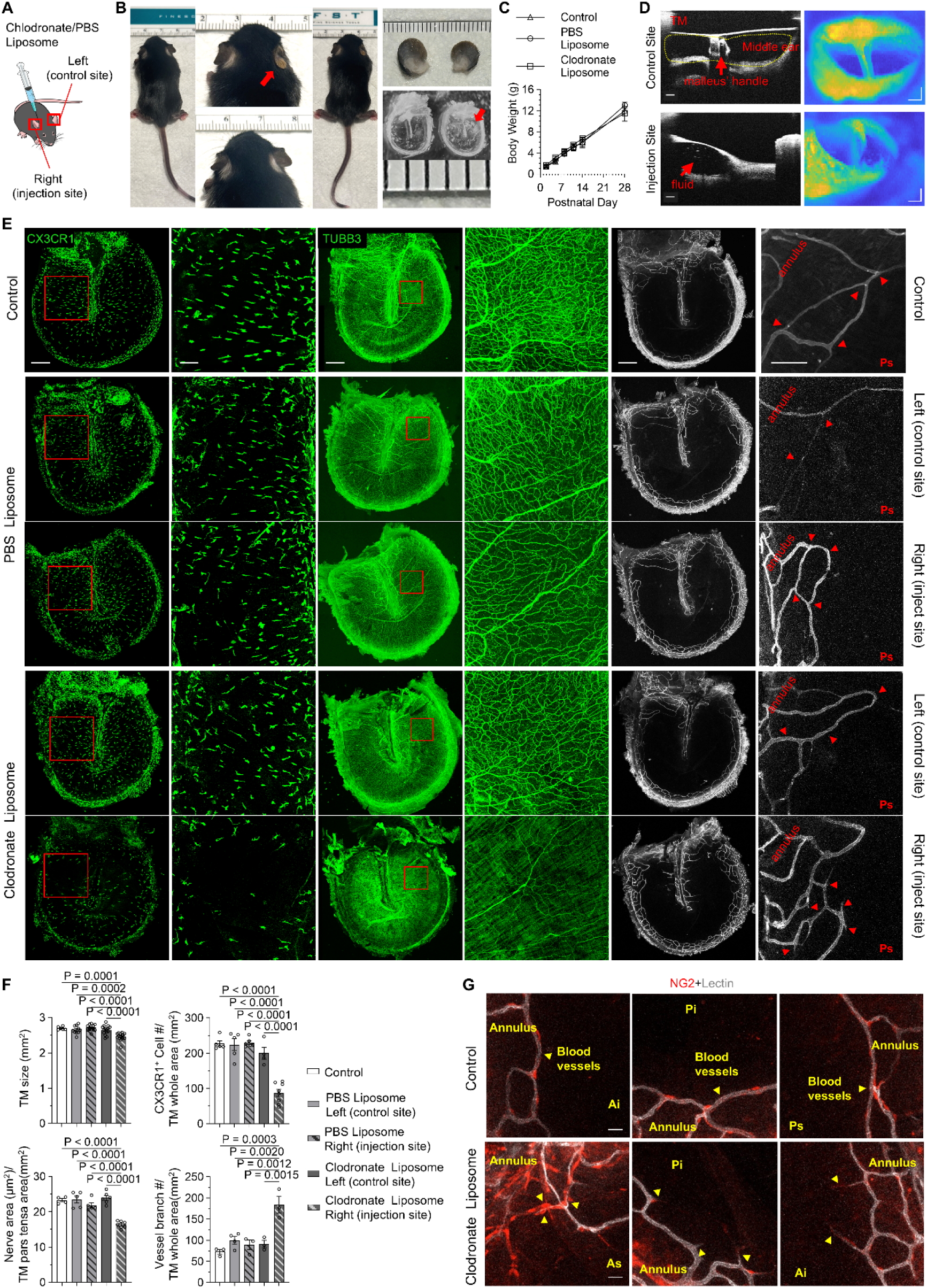
Local macrophage depletion impairs growth and eardrum development. (**A**) Schematic of local macrophage depletion via clodronate liposome injection into the right pinna. (**B**) Representative images showing general animal size (*left*) and pinna/eardrum morphology (*right*) of normal and macrophage-depleted animals at P15. The left arrow indicates that the injected ear fails to stand upright. The right arrows indicate fibrosis on the eardrum at the injection site. Scale bars: 1 mm. (**C**) General animal body weight analysis among different groups (control n=5, PBS liposome n=5, Clodronate liposome n=6). Data were presented as mean ± SEM, statistical significance was calculated through Two-way ANOVA with Šidak’s test (F_Interaction_ (10, 78) =,2.060 F_Raw Factor_ (5, 78) = 561.3, F_Column Factor_ (2, 78) = 2.443). (**D**) *In vivo* 2D OCT B-scans *(left panel)* and En face OCT images *(right panel)* of the eardrum reveal a notable accumulation of fluid in the middle ear and signs of mid-ear infection in the chlodronate group control site (*upper panel)* and injection site (*lower panel*). Scale bar: 150 μm. (**E**) Distribution of macrophages, nerve fibers and blood vessels at P15 in normal, liposome control, and macrophage-depleted animals. Scale bars: 300 μm (*low magnification*), 100 μm (*high magnification*). Arrowheads indicate vessel branching. (**F**) Quantitative analysis of eardrum size (control n=6, PBS liposome left n=10, PBS liposome right n=14, Clodronate liposome left n=15, Clodronate liposome left n=16), macrophage density (control n=5, PBS liposome left n=5, PBS liposome right n=6, Clodronate liposome left n=4, Clodronate liposome left n=8), nerve fiber density (control n=4, PBS liposome left n=5, PBS liposome right n=5, Clodronate liposome left n=6, Clodronate liposome left n=7), and vessel branching(control n=3, PBS liposome left n=4, PBS liposome right n=3, Clodronate liposome left n=3, Clodronate liposome left n=3) density at P15. Data were presented as mean ± SEM, statistical significance was calculated through One-way ANOVA with Tukey’s test (F_TM size_ (4, 56) = 13.18, FCx3Cr1^+^ cell density (4, 23) = 35.67, Fnerve fiber density (4, 22) = 26.76, Fvessel branch density (4, 11) = 13.36, (**G**) Confocal images depicting vessel sprouting and pericyte accumulation in macrophage-depleted versus control mice at P15. Ai: anteroinferior quadrant; Pi: posteroinferior quadrant; Ps: posterior superior quadrant. Scale bar: 25 μm. (**G**) Confocal images depicting vessel sprouting and pericyte accumulation in macrophage-depleted versus control mice at P15. Ai: anteroinferior quadrant; Pi: posteroinferior quadrant; Ps: posterior superior quadrant. Scale bar: 25 μm.

To assess the impact of macrophage depletion on eardrum function, we used a non-invasive optical coherence tomography (OCT) vibrometry system to measure eardrum vibrations in adult mice. OCT imaging revealed middle-ear abnormalities following macrophage depletion. In systemic depletion, middle-ear fluid accumulation and fibrotic changes were frequently observed (Fig. 5F and G), and in local depletion these changes were consistently detected at the injected ear (Fig. 6D). During OCT imaging, we also observed narrowing of the external ear canal in macrophage-depleted ears in both the systemic and local depletion models; this was a qualitative impression rather than a quantified metric. In many affected ears, the presence of fluid/fibrosis coincided with absent or unreliable vibration readouts (Fig. S6B), limiting the interpretability of OCT vibrometry measurements in this setting. Accordingly, we focus on OCT structural imaging as evidence of middle-ear pathology following macrophage depletion. In the systemic depletion group, hearing thresholds were significantly elevated (Fig. S4B). This hearing loss is likely associated with middle-ear fluid or fibrosis (Fig. 5F), and with sensory hair cell loss following macrophage depletion, as previously reported (*31*), although additional systemic effects may also contribute.

## Discussion

This study provides compelling evidence that TRMs are abundantly distributed within the eardrum and undergo substantial changes in both morphology and population dynamics from the neonatal period into adulthood. Along with ultrastructural identification, our data supports that these CX3CR1⁺ cells are bona fide macrophages positioned within the fibrous middle layer of the TM. These macrophages exhibit heterogeneity, arising from multiple origins, with a major embryonic contribution and progressively increasing postnatal input from the GMP lineage during maturation. Coinciding with TRM maturation, we observed the postnatal expansion and structural organization of the eardrum’s vascular and peripheral neural networks. Using advanced 3D imaging, we identified a close spatial and temporal association between TRMs, blood vessels, and nerve fibers. Importantly, neonatal depletion of TRMs resulted in smaller eardrums, immature vasculature, and disorganized nerve fibers during early postnatal development, and was accompanied by frequent middle-ear abnormalities consistent with elevated infection risk, including fluid accumulation and fibrotic changes detected by OCT, highlighting a critical role for TRMs in orchestrating the structural and functional maturation of the eardrum while maintaining middle-ear homeostasis during this vulnerable developmental window.

TRMs stand guard as vigilant sentinels of innate immunity, stationed throughout nearly every tissue in the body (*31–33*). The eardrum is a delicate, multilayered structure that faces continuous environmental challenges, including noise, pressure fluctuations, and invading pathogens. Beyond its essential role in sound conduction and auditory acuity, the eardrum functions as a protective barrier separating the external environment from the middle ear cavity. During the postnatal period—a critical window of tissue adaptation—TRMs positioned within the eardrum are well placed to contribute to local immune defense by surveilling the tissue and responding to bacterial and viral threats. Consistent with this idea, the middle ear is especially susceptible to infection during early childhood, as evidenced by the high incidence of otitis media (*34*) This heightened vulnerability may reflect dynamic remodeling and high density of macrophages in developing tympanic tissue. Notably, depletion of TRMs from postnatal day 2 to 15 increased the incidence of middle-ear abnormalities in mice, including fluid accumulation and fibrosis. In the systemic depletion model, these abnormalities occurred alongside increased mortality and reduced body weight (Fig. 5C-G), consistent with broader systemic effects of macrophage depletion. To minimize systemic confounds, macrophages were also depleted locally using clodronate liposomes. Local depletion produced pronounced tissue-restricted phenotypes, including abnormal pinna morphology and middle-ear abnormalities detected by optical coherence tomography (OCT) at the injected side (Fig. 6D). OCT imaging further revealed an apparent narrowing of the external ear canal in macrophage-depleted ears in both systemic and local depletion models. This observation was based on qualitative OCT imaging rather than quantitative measurement, as imaging angles were not standardized; future studies incorporating post-dissection measurements of the auditory meatus will be necessary to rigorously evaluate this finding. Although more targeted strategies, such as diphtheria toxin-mediated depletion, could further clarify the ear-specific roles of eardrum TRMs in vivo, such approaches are limited by reported neuropathy and ototoxicity(*28*).

Beyond their classical role in immune surveillance, accumulating evidence indicates that TRMs play critical roles in tissue morphogenesis. Notably, TRMs contribute to neurogenesis and vascular remodeling during embryonic and postnatal development. For example, microglia in the central nervous system guide axonal growth, promote myelination, and are essential for synaptic pruning (*35–37*); disruption of neuron-microglia signaling impairs brain connectivity and causes long-term functional deficit (*38*). Likewise, alveolar macrophages contribute to lung development (*39*), early studies have also shown that ocular TRMs regulate vascular regression during eye development (*40*). In our study, TRMs in the eardrum were closely aligned with nascent blood vessels and immature nerve bundles during early postnatal development. Their transcriptional profiling revealed enrichment of genes associated with angiogenesis, axon guidance, and growth factor signaling pathways (Fig. 4G and H), supporting a potential role in coordinating neurovascular maturation. Consistent with this, our scRNA data analysis indicates that TRMs comprise distinct “M1-like” and “M2-like” transcriptional programs, as shown in Fig. 4E, which suggest functional specialization. M1-like macrophages are found to be enriched for neurovascular growth and remodeling processes, such as Schwann cell proliferation, axon regeneration, sprouting angiogenesis, and VEGF regulation. In contrast, M2-like macrophages are enriched for synapse pruning and axon-injury response, while also retaining angiogenesis-related programs. Clearly, both macrophage types exist along a spectrum and respond to microenvironmental cues in neurovascular development of the eardrum, in addition to their defense roles, which are not excluded in this study.

Our depletion studies further demonstrate the essential roles of these cells in the coordinated maturation of eardrum structure and its neurovascular networks. Systemic TRM depletion led to smaller eardrums, reduced vascular branching, decreased nerve fiber density (Fig. 5E, H and I), abnormal pericyte accumulation, and persistent vascular sprouting (Fig. 5J). Regional TRM depletion produced similar effects, including smaller pinna and more pronounced abnormalities in pericyte distribution (Fig. 6B-G). The eardrum’s vascular network contains abundant pericytes, which are critical for vascular stabilization and maturation (*26*). In contrast, its neural network consists of densely packed fibers without neuronal cell bodies. To investigate intercellular communication, we collected only macrophage and pericyte cell types for scRNA-seq, given the absence of neuronal cell bodies in the eardrum neural network. CellChat analysis of scRNA-seq data identified potential signaling interactions between TRMs and pericytes, particularly involving Oncostatin M (Osm), a cytokine implicated in vascular remodeling (Fig. 4I). While the molecular mechanisms underlying this pathway remain uncharacterized, TRM depletion was associated with abnormal pericyte morphology (Fig. 5J), suggesting a disruption in TRM-pericyte communication. Future studies are required to elucidate the molecular basis of this interaction.

The eardrum is a specialized structure composed of both epithelial and connective tissues, endowing it with unique vibratory properties distinct from those of skin or mucosa. Abnormal eardrum development can result in conductive hearing loss, leading to delays in speech, language acquisition, and cognitive development (*41*). In our study, OCT structural imaging revealed frequent middle-ear fluid accumulation and fibrotic changes following macrophage depletion (Fig. 5F and G; Fig. 6D). In many affected ears, these abnormalities coincided with absent or unreliable vibration readouts (Fig. S6B), limiting interpretation of OCT vibrometry in this setting; therefore, we emphasize OCT structural imaging as evidence of middle-ear pathology. These structural abnormalities are consistent with mechanical and pressure-related contributors to conductive impairment(*42*). Additionally, we would like to highlight our previous work which shows that the eardrum is heavily innervated by TRPV1-positive sensory fibers (*2*). The TRPV channel is known to respond to mechanical and thermal stimuli (*43*). Macrophage depletion results in sensory nerve impairment, which may also contribute to the observed reduction in vibration sensitivity. In the systemic depletion cohort only, ABR thresholds were elevated (Fig. S4B), consistent with hearing impairment in vivo. Because systemic clodronate liposome treatment affects macrophages beyond the eardrum (including cochlear macrophages; Fig. S4C), these functional changes likely reflect a combination of conductive contributions from middle-ear pathology and additional systemic/inner-ear effects that cannot be excluded. In addition, macrophage depletion has been reported to be associated with auditory functional impairments, including mistargeted spiral ganglion neuron (SGN) axons and retracted or excessively pruned peripheral nerve fibers (*31*), which may further contribute to elevated ABR thresholds. Additionally, we observed systemic growth retardation post-macrophage depletion, indicated by smaller body size and lower survival rates, aligning with macrophages’ roles in energy metabolism and organ development (*44*). These findings suggest that macrophage deficiency can cause localized and systemic developmental impairments. A tissue-restricted depletion model would be ideal for focusing on TRMs in the eardrum and will be important in future studies to isolate TM-intrinsic TRM mechanisms. Nonetheless, our results highlight a critical role for postnatal TRMs in coordinating neurovascular maturation of the eardrum and maintaining middle-ear homeostasis during a vulnerable developmental window. Furthermore, the results may also have clinical relevance, as eardrum abnormalities have long been observed in humans (*15, 16*), although the underlying mechanisms remain unclear. This study offers new insight into how the innate immune system influences TM structure and function.

This study has several limitations. First, clodronate liposome treatment induces systemic macrophage depletion, so middle-ear abnormalities and growth phenotypes may reflect broader immune suppression rather than TM-restricted effects. Although local pinna injections largely spared overall growth and still produced ear-specific abnormalities, a tissue-restricted TRM depletion strategy will be needed to isolate TM-intrinsic mechanisms. Second, although OCT structural imaging robustly detected fluid and fibrotic changes, vibration readouts were often absent or unreliable in affected ears, limiting interpretation of OCT vibrometry under these conditions. Third, the observed narrowing of the external ear canal was a qualitative impression rather than a quantified metric. Finally, CellChat predicts ligand–receptor interactions but does not establish causality; future studies will be required to test specific TRM-pericyte signaling mechanisms (including Osm) using targeted perturbations and tissue-restricted depletion approaches.

Nonetheless, our research for the first time demonstrates that postnatal TRMs coordinate neurovascular growth of the eardrum and support middle-ear balance during a key developmental period. These findings may have important clinical implications, as eardrum abnormalities and children’s susceptibility to middle-ear infections are well documented (*15, 16*), but the underlying mechanisms are not fully understood. Our work provides a new perspective on the factors shaping eardrum structure and function.

### Materials and Methods Experimental Design

The objective of this study was to define the distribution, developmental origin, transcriptional programs, and functional roles of tissue-resident macrophages (TRMs) in the tympanic membrane (TM) during postnatal development. We combined (i) reporter-based whole-mount imaging across postnatal timepoints, (ii) Cre-lox fate mapping of embryonic and postnatal myeloid lineages, (iii) ultrastructural identification by TEM with HRP uptake, (iv) scRNA-seq of TM macrophages and pericytes at P15 with GO enrichment and CellChat inference, and (v) systemic versus local clodronate-liposome depletion to test functional contributions to TM growth, neurovascular maturation, and middle-ear abnormalities assessed by confocal imaging, OCT structural imaging, and ABR. Animals were allocated to experimental groups by litter availability and genotype; imaging and quantification were performed in a blind manner when stated below. Sample sizes (n) and statistical tests are reported in figure legends.

### Animal models

The following mouse strains were used in this study: B6.129P2(Cg)-*Cx3cr1^tm1Litt^*/J (*CX3CR1^EGFP^*, JAX #005582), B6.129P2(Cg)-*Cx3cr1^tm2.1(cre/ERT2)Litt^*/WganJ (*CX3CR1^CreER^*, JAX #021160), C57BL/6J-*Ms4a3^em2(cre)Fgnx^*/J (*Ms4a3^cre^*, JAX #036382), B6.Cg-*Gt(ROSA)26Sor*^tm6*(CAG-ZsGreen1)Hze*^/J (R26*^ZsGreen^*, JAX # 007906), B6.Cg-Gt(ROSA)26Sor*^tm9(CAG-tdTomato)Hze^*/J (R26*^tdTomato^*, JAX #007909), , B6.Cg-Tg(Csf1r-EGFP)1Hume/J (*CSF1R^EGFP^*, JAX #018549), Tg(Cspg4-DsRed.T1)1Akik/J (*NG2^DsRed^*, JAX #008241), B6.129-*Trpv1^tm1(cre)Bbm^*/J (*TRPV1^Cre^*, strain # 017769). Additional strain, FVB-Tg(Csf1r-cre/Esr1*)1Jwp/J (*Csf1r-Mer-iCre-Mer)*, was generously provided by Dr. Benjamin A. Alman (Duke University, Durham, NC). All transgenic mice were maintained through in-lab breeding and routinely genotyped for validation. Both male and female mice were used for experiments.

*CX3CR1^EGFP^* mice were utilized at postnatal day 1 (P1) through 30 to assess steady-state macrophage distributions in the eardrum. For fate mapping experiments investigating macrophages ontogeny, *R26^tdTomato^*mice were crossed with *Csf1r-Mer-iCre-Mer*, *CX3CR1^CreER^*, and *Ms4a3^cre^* mice. Cre-mediated recombination was induced at embryonic days (E) 8.5 or E12.5. Embryonic development stage was defined relative to the detection of a vaginal plug, designated as E0.5 (Fig. 2A). To induce recombination, pregnant dams were administered intraperitoneal injections of 4-hydroxytamoxifen (4-OHT, Sigma Aldrich, Cat# H6278) at a dose of 75 μg/g body weight, supplemented with 37.5 μg/g of corn oil (Sigma Aldrich, Cat# C8267) to mitigate the risk of fetal abortion. All animal experiments reported were approved by the Oregon Health & Science University Institutional Animal Care and Use Committee (IACUC IP00000968).

### Eardrum immunostaining

The auditory bulla and capsule, containing the eardrum and middle ear, were isolated and fixed in 4% paraformaldehyde (PFA) overnight at 4°C. Following fixation, tissues were washed three times with 1× PBS and the eardrum was carefully dissected. For F4/80 (Invitrogen, Cat# 14480185) labeling, samples were incubated in a blocking/permeabilization solution consisting of 0.25% Triton X-100 and 10% goat serum (GS) in 1× PBS for 1 hour at room temperature (RT). Tissues were then transferred into the primary antibody solution, diluted in the same blocking/permeabilization buffer, and incubated overnight at 4°C. For nerve fiber labeling using β-III tubulin (TUBB3, Abcam, Cat# ab78078), eardrums were first permeabilized in 1% Triton X-100 for 2 hours at RT, followed by blocking in 10% GS and 1% bovine serum albumin (BSA) in 1× PBS for 1 hour. Samples were then incubated overnight at 4°C with the primary antibody against TUBB3, diluted in 10% GS and 1% BSA.

After primary antibody incubation, all samples were washed three times with 1× PBS and then incubated with fluorescence-conjugated secondary antibodies (diluted in blocking/permeabilization buffer) for 1 hour at RT. Samples were again washed three times with PBS, mounted on glass slides using Antifade Mounting Medium with DAPI (Thermo Fisher Scientific), and imaged under a 10× objective using an Olympus FV1000 laser-scanning confocal microscope (Olympus, Japan). Specifically, we utilized Z-stack (3 μm per step, ∼ 30 steps) imaging to capture the full thickness of the TM, thereby encompassing all the layers of the eardrum. Due to the conical shape and large surface area of the eardrum, capturing the entire structure in a single field of view was not feasible. Therefore, the TM was imaged in four distinct regions, and the resulting images were manually stitched for full coverage. Laser intensity, step size, and resolution were kept consistent across all scans to ensure comparability. In addition, selected samples were imaged using a Zeiss LSM980 confocal microscope (Zeiss, Germany), also under a 10× objective, utilizing the Z-stack and tile stitching tools to acquire high-resolution, full-field images of the eardrum.

### Transmission electron microscopy (TEM)

To identify macrophages, mice were intraperitoneally injected with a solution of horseradish peroxidase type II (HRP type II, Sigma-Aldrich, USA) at a concentration of 15 mg/ml (100 μL) 18 hours prior to sacrifice, as previously reported (Shi, 2010). The eardrum was then isolated and fixed in a solution containing 2.5% glutaraldehyde and 2% paraformaldehyde in phosphate buffer (0.1 M, pH 7.4, 250-300 mL per animal). The samples were sent to the Multiscale Microscopy Core at OHSU for further processing. Briefly, the tissues were post-fixed in 1% osmium (Electron Microscopy Sciences, Hatfield, PA), dehydrated through a graded series of alcohol, and embedded in Embed 812 (Electron Microscopy Sciences, Hatfield, PA). They were then sectioned, stained with lead citrate and uranyl acetate (both from Electron Microscopy Sciences, Hatfield, PA), and viewed using a Philips CM 100 transmission electron microscope (Philips/FEI Corporation, Eindhoven, Holland).

### Macrophage quantification

Macrophage quantification and measurements of the eardrum area were performed using Fiji (ImageJ, version 1.54p). Macrophage density was calculated as the number of CX3CR1⁺ cells per 10⁶ μm² by dividing the total number of CX3CR1⁺ cells by the measured area of the eardrum. To quantify CX3CR1⁺ cells from postnatal day 1 (P1) to postnatal day 30 (P30), 3D confocal image stacks were acquired using an Olympus confocal microscope. Final 2D representations of the TM were created by stitching and projecting these image stacks. Each image was individually masked and thresholded, and CX3CR1⁺ cells were quantified using the “Analyze Particles” function in Fiji (Fig. S7). A size filter ranging from 50 to 400 pixels² was applied to identify macrophages while excluding background noise and debris. Cells located at the edges of the images were excluded from the analysis to avoid partial counting artifacts. All quantifications were performed in a blind manner to minimize bias.

### Single-cell RNA sequencing and annotation

*CX3CR1^EGFP^*; *NG2^DsRed^*mice at P15 were used for scRNA-seq. Temporal bones were harvested from 10 mice (20 capsules) and placed in ice-cold RPMI medium. Eardrums were dissected and transferred to 1.5 mL microcentrifuge tubes containing 1 mL of RPMI medium supplemented with 1.5 mg/mL collagenase IV (Gibco, Cat# 17104019) and 1 mg/mL DNase I (Roche, Cat# 10104159001). Tissues were minced with sterile micro scissors and digested at 37°C for 30 minutes with gentle shaking (100 rpm). Following digestion, the mixture was centrifuged at 300 × g for 5 minutes at 4°C. The pellet was further dissociated in TrypLE Express (Gibco, Cat# 12604013) at 37°C for 10 minutes, with pipette trituration every 5 minutes using a 1000 μL tip to generate a single-cell suspension. The suspension was passed through a 70 μm cell strainer (Falcon), washed once with PBS, and resuspended in 0.04% BSA in PBS prior to fluorescence-activated cell sorting (FACS). Sorted cells were submitted to the Massively Parallel Sequencing Shared Resource (MPSSR) at Oregon Health & Science University (OHSU) for scRNA-seq. Prior to library preparation, cells were assessed for viability and concentration using Trypan blue staining and the Bio-Rad TC20 Automated Cell Counter. Only samples with > 85% viable cells were processed. Cells were loaded onto a 10x Genomics Chromium chip, targeting a recovery of at least 2,500 single cells.

Raw sequencing data generated by 10x Genomics were processed using the Seurat package (v5.1.0) (*45*) in R. Quality control filters retained cells with: >200 and <6000 detected genes, 200 – 50,000 UMI counts, and <5% mitochondrial gene content. Ribosomal content was also calculated (percent.ribo; Rps/Rpl genes) and cells with percent.ribo <40% were retained (Supplementary Fig. 4a-b). Gene expression data were normalized using the Log Normalize method (scale factor = 10,000), and highly variable features were identified. Data were then scaled across all genes. Principal component analysis (PCA) was performed on variable genes, and significant components were selected based on elbow plot inspection. Neighbor graphs were constructed using the first 10 PCs, clustering was performed using the Louvain algorithm (resolution = 0.5), and UMAP was computed using the selected PCs. Cluster identities were manually annotated using canonical marker genes. Doublets/multiplets were identified using scDblFinder after conversion to a SingleCellExperiment object, and only singlets were retained for downstream analyses (Fig. S4C) (*46*). After singlet filtering, cells were gated to enrich macrophages and pericytes by retaining cells expressing either *Cx3cr1* (>0) or the pericyte marker *Cspg4* (>0), while excluding Itgax⁺ (*Cd11c*⁺) Dendritic/Langerhans cells (*Itgax* = 0). These cells, along with identified pericytes, were subset and re-clustered for comparative analysis (Fig. S4D). Macrophage subtype markers were identified using FindAllMarkers (Wilcoxon test; only.pos = TRUE; log fold change threshold >0.25; min.pct ≥0.10). To reduce technical and non-informative signals, mitochondrial/ribosomal/histone genes and common housekeeping genes were removed from marker lists prior to enrichment analyses. For GO Biological Process enrichment, the top 500 markers per macrophage subtype were exported and submitted for downstream GO analysis on PANTHER (*47–49*). Finally, CX3CR1^+^ macrophages and pericytes were analyzed using CellChat (v2.1.2) (*50*) to infer ligand–receptor-mediated intercellular communication. Macrophage and pericyte Seurat objects were merged, the RNA assay layers were joined (JoinLayers), and cell identities were set to M1-like, M2-like, and PC before CellChat inference. CellChat analysis used the mouse ligand–receptor database (CellChatDB.mouse) and included subsetData, identification of overexpressed genes/interactions, computation of communication probability and pathway-level communication, network aggregation, and visualization of signaling networks (circle plots and bubble plots).

### Macrophage depletion with clodronate liposomes

For systemic depletion, Clodronate liposomes (Liposome Clodronate, Netherlands) were used to deplete macrophages according to the manufacturer’s instructions. A suspension of clodronate liposomes was administered at a dose of 10 μL per gram of body weight. The first dose was delivered to P2 mice via intravenous (*i.v*.) injection, followed by intraperitoneal (*i.p*.) injections every three days until P15 to maintain macrophage depletion throughout early postnatal development. To trace the kinetics of liposomes in eardrum TRMs, DiI-labeled liposomes (Liposome Clodronate, Netherlands) were administered at the same dosage to P2 mice. Eardrums were collected at 24-, 48-, and 72-hour post-administration for analysis (Fig. S4A).

For local depletion, Clodronate liposomes (Liposome Clodronate, Netherlands) were administered by subcutaneous *(s.c.)* injection into the right pinna at a dose of 5 μL per gram of body weight. The first dose was delivered to P2 mice, followed by repeat injections every three days until P15 to maintain macrophage depletion throughout early postnatal development.

### Assessment of vascular area, branching, vessel diameter, and nerve fiber density

To visualize blood vessels, mice were anesthetized and injected with Lectin-DyLight 649 (DL-1178, Vector laboratories). The dye was diluted in 0.1 M PBS to a concentration of 20 μg/mL (100 μL total volume) and administered via retro-orbital sinus injection 10 minutes prior to sacrifice(*51*). Nerve fibers were visualized in whole-mount eardrum preparations using either TUBB3 immunolabeling, or endogenous TRPV1 fluorescence in *TRPV1^Cre^; R26^tdTomato^* mice. We previously validated that TRPV1^+^ signals closely correlate with β-III tubulin-labeled nerve fibers in the eardrum(*2*). Quantification of blood vessel area, nerve fiber area, vessel branch number, and eardrum areas were performed using Fiji (ImageJ, v1.51t). Vessel diameters were assessed by measuring the width of 10 randomly selected vessels per sample within a defined region of interest. For vessel branching analysis, images were skeletonized and branching points were quantified using the “Analyze Skeleton (2D/3D)” plugin in ImageJ. Vascular density was defined as vascular 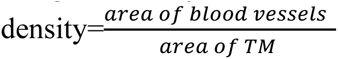 Vessel branch density was defined as vascular 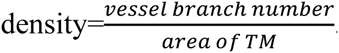 Nerve fiber density was defined as nerve fiber density= 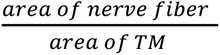

### Eardrum mechanical function measurement using optical coherence tomography (OCT)

To assess eardrum vibratory function, control, PBS liposome, and clodronate liposome-treated mice were anesthetized with ketamine (100 mg/kg) and xylazine (10 mg/kg). Supplemental doses were administered as needed to maintain areflexia. The left pinna and external auditory canal were removed, and tissue surrounding the eardrum was bluntly dissected to maximize the visual field. While the bony ear canal limited full visualization of the TM, a wide field of view was obtained.

The animal was positioned on its flank on a heating pad and secured with a head holder. To minimize respiratory motion, a stabilizing loop fashioned from a paperclip and mounted on a copper rod was placed around the TM using a micromanipulator. An 8 kHz sound stimulus was delivered using a modified Sony MDR-V6 headphone speaker, positioned 30 cm from the animal. A Piezotronics microphone, powered by a 480C02 ICP signal conditioner, was placed 3 mm from the ear canal, facing the speaker. The TM was visualized using a Thorlabs Telesto III OCT microscope (central wavelength: 1300 nm), equipped with an LSM03 5× objective. A ∼2 mm scan line, positioned 1 mm parallel to the pars tensa/pars flaccida interface, was used. The OCT microscope focus was adjusted to optimize OCT signal strength from the eardrum while minimizing autocorrelation artifacts.

Once the sound system and OCT microscope were aligned, a continuous 8 kHz raised cosine tone was generated using a Stanford SR830 Lock-in amplifier. The speaker output was calibrated to ensure 81 dB SPL output. The stimulus was delivered via a National Instruments board, synchronized with custom OCT acquisition software (10,000 samples; 76 kHz sampling rate; 135 ms stimulus duration). Structural OCT scans were acquired in both 2D and 3D modes (X-axis: 512 A-lines across 2 mm (averaged 10 times), Y-axis: 100 A-lines across 2 mm, Z resolution: ∼3 μm).

Vibration data were analyzed using custom MATLAB software: two successive fast Fourier transforms were conducted, the first giving the morphological structure, and the second providing TM motion via processing of the phase data (*52, 53*). Using morphological OCT images, the TM was selected in a region of interest. Pixels satisfying a 6 dB SNR criterion were extracted, and the displacement was expressed as the average of the responsive TM pixels along the Z axis of the scan. Displacement data are expressed in absolute values (nm).

### Auditory brainstem response (ABR)

ABR audiometry was performed at P21 to assess hearing function, as previously described (*51*). Mice were anesthetized with ketamine and xylazine and placed on a heating pad maintained at 37°C inside a sound-isolated chamber. Subdermal needle electrodes were inserted near the test ear (active), at the vertex (reference), and on the contralateral ear (ground). Each ear was stimulated independently using a closed-tube sound delivery system sealed into the ear canal. Tone bursts (1 ms rise time) at 8, 16, 24, and 32 kHz were presented, and the resulting auditory brainstem responses were recorded. Hearing thresholds were determined for each frequency and for each ear based on the lowest stimulus intensity that elicited a reproducible response.

### Statistical Analysis

All statistical analyses were performed using GraphPad Prism (version 10.4.1). For comparisons involving three or more groups, one-way ANOVA followed by Tukey’s multiple comparisons test was used. For body weight and ABR threshold comparisons across frequencies, two-way ANOVA followed by Šídák’s multiple comparisons test was used. A p-value < 0.05 was considered statistically significant. All data are presented as mean ± SEM. Values for N, P, and the specific statistical test performed for each experiment are reported in the appropriate figure legend and/or main text.

## Acknowledgments

We would like to express our gratitude to MPSSR at OHSU for helping with the collection of our scRNA-seq dataset, and Dr. Anders Fridberger for the development of OCT data acquisition software.

## Author contributions

Conceptualization: XRS

Methodology: YZ, PW, LN, GB, KS

Investigation: YZ, PW, LN, GB

Formal analysis: YZ, PW

Validation: YZ, PW

Visualization: YZ, PW, LN, GB, KS

Supervision: XRS

Writing—original draft: YZ, PW, GB, XRS

Writing—review & editing: YZ, PW, AK, GB, XRS

## Data and materials availability

All data are available in the main text or the supplementary materials. Source data are provided with this paper. Data supporting the findings of this study are available in the article, its Supplementary Information, the source data file, and from the corresponding author upon request. Single-cell RNA-seq data will be deposited at GEO: GSE296504 and publicly available as of the date of publication.

## References

1. J. Zhao, I. Andreev, H. M. Silva, Resident tissue macrophages: Key coordinators of tissue homeostasis beyond immunity. Sci Immunol 9, eadd1967 (2024).

2. Y. Zhang et al., Monocyte-derived macrophage recruitment mediated by TRPV1 is required for eardrum wound healing. bioRxiv, (2025).

3. G. Hoeffel, F. Ginhoux, Fetal monocytes and the origins of tissue-resident macrophages. Cell Immunol 330, 5–15 (2018).

4. T. Lazarov, S. Juarez-Carreno, N. Cox, F. Geissmann, Physiology and diseases of tissue-resident macrophages. Nature 618, 698–707 (2023).

5. L. van de Laar et al., Yolk Sac Macrophages, Fetal Liver, and Adult Monocytes Can Colonize an Empty Niche and Develop into Functional Tissue-Resident Macrophages. Immunity 44, 755–768 (2016).

6. S. G. Utz et al., Early Fate Defines Microglia and Non-parenchymal Brain Macrophage Development. Cell 181, 557–573 e518 (2020).

7. S. Y. Tan, M. A. Krasnow, Developmental origin of lung macrophage diversity. Development 143, 1318–1327 (2016).

8. T. N. Shaw et al., Tissue-resident macrophages in the intestine are long lived and defined by Tim-4 and CD4 expression. J Exp Med 215, 1507–1518 (2018).

9. T. A. Wynn, A. Chawla, J. W. Pollard, Macrophage biology in development, homeostasis and disease. Nature 496, 445–455 (2013).

10. D. M. Mosser, K. Hamidzadeh, R. Goncalves, Macrophages and the maintenance of homeostasis. Cell Mol Immunol 18, 579–587 (2021).

11. S. Watanabe, M. Alexander, A. V. Misharin, G. R. S. Budinger, The role of macrophages in the resolution of inflammation. J Clin Invest 129, 2619–2628 (2019).

12. M. J. Mason, Structure and function of the mammalian middle ear. II: Inferring function from structure. J Anat 228, 300–312 (2016).

13. J. P. Fay, S. Puria, C. R. Steele, The discordant eardrum. Proc Natl Acad Sci U S A 103, 19743–19748 (2006).

14. L. Widemar, S. Hellstrom, M. Schultzberg, L. E. Stenfors, Autonomic innervation of the tympanic membrane. An immunocytochemical and histofluorescence study. Acta Otolaryngol 100, 58–65 (1985).

15. G. R. Holt, T. M. Watkins, M. G. Yoder, Assessment of tympanometry abnormalities of the tympanic membrane. Am J Otolaryngol 3, 112–116 (1982).

16. N. W. Todd, Association of abnormal appearance of tympanic membrane with minimal temporal bone pneumatization. A cadaver study. ORL J Otorhinolaryngol Relat Spec 49, 133–137 (1987).

17. B. Chen, R. Li, A. Kubota, L. Alex, N. G. Frangogiannis, Identification of macrophages in normal and injured mouse tissues using reporter lines and antibodies. Sci Rep 12, 4542 (2022).

18. M. Burgess, K. Wicks, M. Gardasevic, K. A. Mace, Cx3CR1 Expression Identifies Distinct Macrophage Populations That Contribute Differentially to Inflammation and Repair. Immunohorizons 3, 262–273 (2019).

19. A. Dos Anjos Cassado, F4/80 as a Major Macrophage Marker: The Case of the Peritoneum and Spleen. Results Probl Cell Differ 62, 161–179 (2017).

20. D. J. Lim, Tympanic membrane. Electron microscopic observation. I: pars tensa. Acta Otolaryngol 66, 181–198 (1968).

21. A. Szymanski, J. Toth, M. Ogorevc, Z. Geiger, in StatPearls. (Treasure Island (FL), 2025).

22. Y. Tian et al., The structural characteristics of mononuclear-macrophage membrane observed by atomic force microscopy. J Struct Biol 206, 314–321 (2019).

23. N. Means, C. K. Elechalawar, W. R. Chen, R. Bhattacharya, P. Mukherjee, Revealing macropinocytosis using nanoparticles. Mol Aspects Med 83, 100993 (2022).

24. X. Shi, Resident macrophages in the cochlear blood-labyrinth barrier and their renewal via migration of bone-marrow-derived cells. Cell Tissue Res 342, 21–30 (2010).

25. Z. Liu et al., Fate Mapping via Ms4a3-Expression History Traces Monocyte-Derived Cells. Cell 178, 1509–1525 e1519 (2019).

26. X. Shi, Research advances in cochlear pericytes and hearing loss. Hear Res 438, 108877 (2023).

27. T. Kubin et al., The Role of Oncostatin M and Its Receptor Complexes in Cardiomyocyte Protection, Regeneration, and Failure. Int J Mol Sci 23, (2022).

28. H. Konishi et al., Exposure to diphtheria toxin during the juvenile period impairs both inner and outer hair cells in C57BL/6 mice. Neuroscience 351, 15–23 (2017).

29. A. Goldwich, A. Steinkasserer, A. Gessner, K. Amann, Impairment of podocyte function by diphtheria toxin--a new reversible proteinuria model in mice. Lab Invest 92, 1674–1685 (2012).

30. L. Mann et al., CD11c.DTR mice develop a fatal fulminant myocarditis after local or systemic treatment with diphtheria toxin. Eur J Immunol 46, 2028–2042 (2016).

31. L. N. Brown et al., Macrophage-Mediated Glial Cell Elimination in the Postnatal Mouse Cochlea. Front Mol Neurosci 10, 407 (2017).

32. G. Hu, J. W. Christman, Editorial: Alveolar Macrophages in Lung Inflammation and Resolution. Front Immunol 10, 2275 (2019).

33. P. Chiaranunt, S. L. Tai, L. Ngai, A. Mortha, Beyond Immunity: Underappreciated Functions of Intestinal Macrophages. Front Immunol 12, 749708 (2021).

34. A. A. Altamimi et al., Recurrent otitis media and behaviour problems in middle childhood: A longitudinal cohort study. J Paediatr Child Health 60, 12–17 (2024).

35. N. B. McNamara et al., Microglia regulate central nervous system myelin growth and integrity. Nature 613, 120–129 (2023).

36. Y. Fujita, T. Yamashita, Mechanisms and significance of microglia-axon interactions in physiological and pathophysiological conditions. Cell Mol Life Sci 78, 3907–3919 (2021).

37. M. Colonna, O. Butovsky, Microglia Function in the Central Nervous System During Health and Neurodegeneration. Annu Rev Immunol 35, 441–468 (2017).

38. Y. Zhan et al., Deficient neuron-microglia signaling results in impaired functional brain connectivity and social behavior. Nat Neurosci 17, 400–406 (2014).

39. A. Kulle, A. Thanabalasuriar, T. S. Cohen, M. Szydlowska, Resident macrophages of the lung and liver: The guardians of our tissues. Front Immunol 13, 1029085 (2022).

40. R. A. Lang, J. M. Bishop, Macrophages are required for cell death and tissue remodeling in the developing mouse eye. Cell 74, 453–462 (1993).

41. T. Sooriyamoorthy, O. De Jesus, Conductive hearing loss. (2020).

42. S. D. Esteves, A. P. Silva, M. B. Coutinho, J. M. Abrunhosa, C. Almeida e Sousa, Congenital defects of the middle ear--uncommon cause of pediatric hearing loss. Braz J Otorhinolaryngol 80, 251–256 (2014).

43. W. Liedtke, Transient receptor potential vanilloid channels functioning in transduction of osmotic stimuli. J Endocrinol 191, 515–523 (2006).

44. F. Guan et al., Tissue macrophages: origin, heterogenity, biological functions, diseases and therapeutic targets. Signal Transduct Target Ther 10, 93 (2025).

45. Y. Hao et al., Dictionary learning for integrative, multimodal and scalable single-cell analysis. Nat Biotechnol 42, 293–304 (2024).

46. P. L. Germain, A. Lun, C. Garcia Meixide, W. Macnair, M. D. Robinson, Doublet identification in single-cell sequencing data using scDblFinder. F1000Res 10, 979 (2021).

47. H. Mi et al., Protocol Update for large-scale genome and gene function analysis with the PANTHER classification system (v.14.0). Nat Protoc 14, 703–721 (2019).

48. H. Mi, P. Thomas, PANTHER pathway: an ontology-based pathway database coupled with data analysis tools. Methods Mol Biol 563, 123–140 (2009).

49. P. D. Thomas et al., PANTHER: Making genome-scale phylogenetics accessible to all. Protein Sci 31, 8–22 (2022).

50. S. Jin, M. V. Plikus, Q. Nie, CellChat for systematic analysis of cell-cell communication from single-cell transcriptomics. Nat Protoc 20, 180–219 (2025).

51. J. Zhang, et al., VEGFA165 gene therapy ameliorates blood-labyrinth barrier breakdown and hearing loss. JCI Insight 6, (2021).

52. R. K. Wang, A. L. Nuttall, Phase-sensitive optical coherence tomography imaging of the tissue motion within the organ of Corti at a subnanometer scale: a preliminary study. J Biomed Opt 15, 056005 (2010).

53. G. Burwood, P. Hakizimana, A. L. Nuttall, A. Fridberger, Best frequencies and temporal delays are similar across the low-frequency regions of the guinea pig cochlea. Sci Adv 8, eabq2773 (2022).

