## Supplemental Figures for "Defects in tissue-resident macrophages lead to smaller eardrums and abnormal neurovascular networks with increased middle-ear infection"

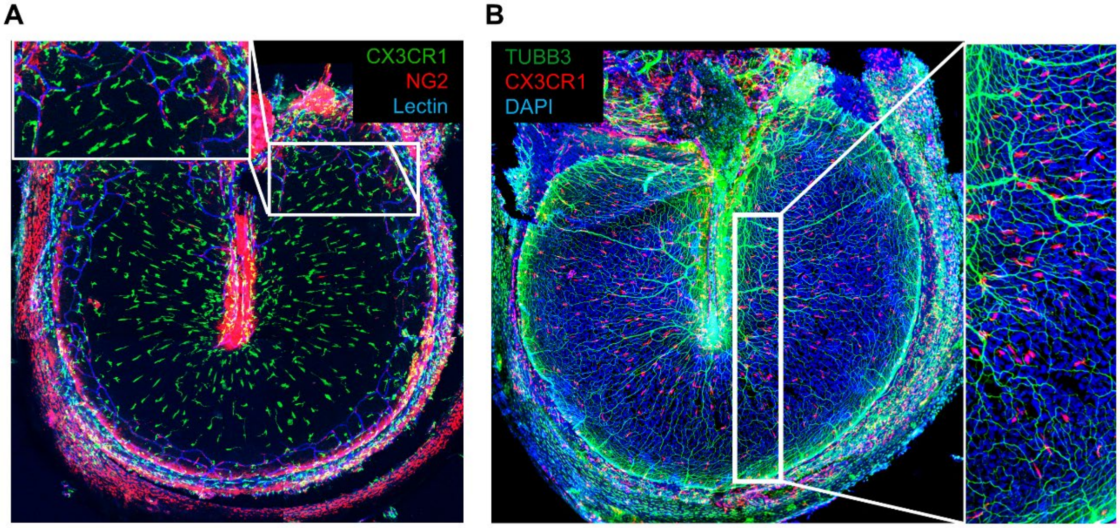


**Fig. S1. Representative confocal images of eardrums collected from P15 mice.** (**A**) representative confocal image of eardrum collected from P15 *CX3CR1*$^{EGFP}$; *NG2*$^{DsRed}$ mice demonstrating the distribution of macrophages (green), pericytes (red) and blood vessels (blue) within the eardrum. (**B**) representative confocal image of eardrum collected from P15 *CX3CR1*$^{EGFP}$ mice, showing close localization of macrophages (red) and nerve fibers (TUBB3, green) within the eardrum. Scale bars, 150 μm.

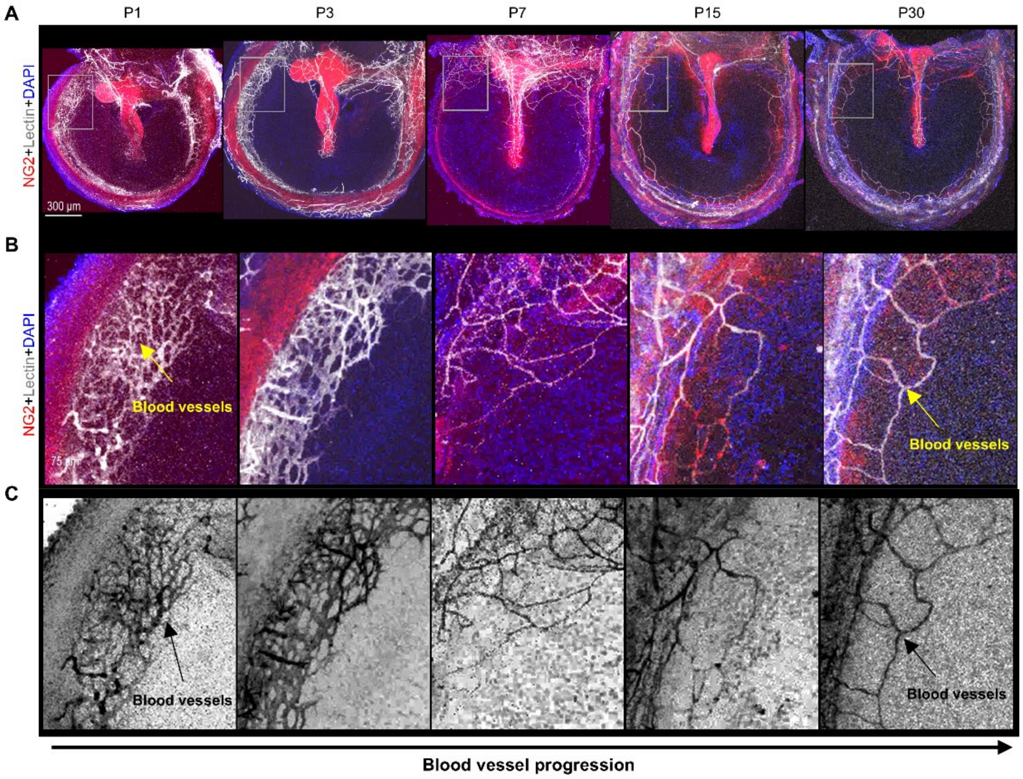


**Fig. S2: Postnatal development of blood vessels in the eardrum**. (**A, B**) Confocal images taken at high and low magnification demonstrate the development of the vascular network in the eardrum of postnatal NG2DsRed mice. (**C**) Inverted images created using Adobe Photoshop (version X6) further show the progression of vascular development. Scale bars: 300 μm (**A**).

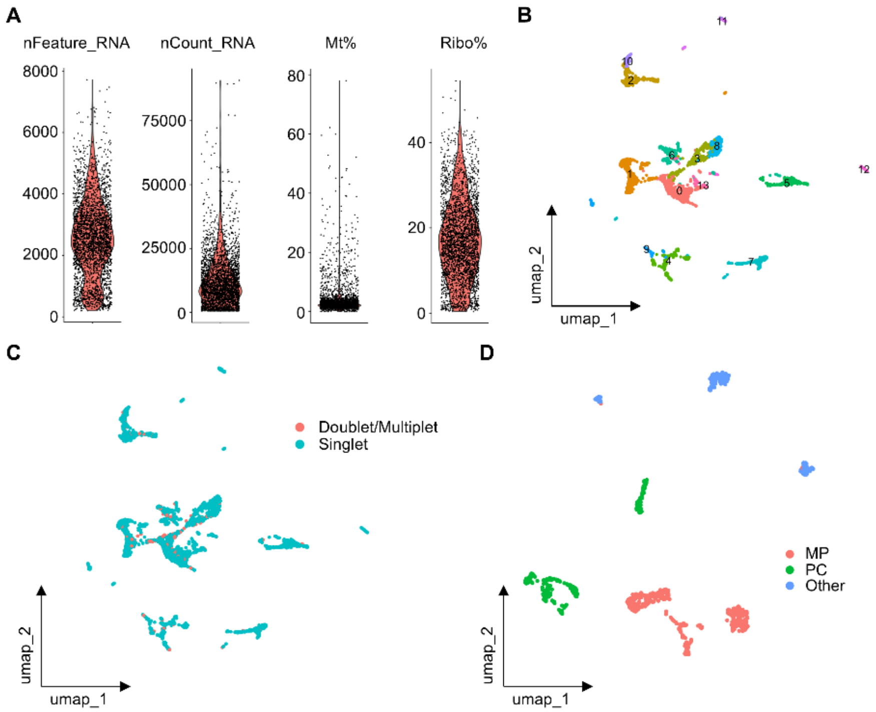


**Fig. S3**. **Quality control and clustering of TM macrophages (MPs) and pericytes (PCs).** (**A**) Initial QC: violin plots of nFeature_RNA, nCount_RNA, and mitochondrial (mt)/ribosomal (ribo) content with applied thresholds. (**B**) UMAP embedding after initial QC. (**C**) UMAP highlighting putative doublets and singlets following scDblFinder doublet detection. (**D**) UMAP of Mps, PCs, and other cells after singlet filtering and MP/PC canonical-marker gating.

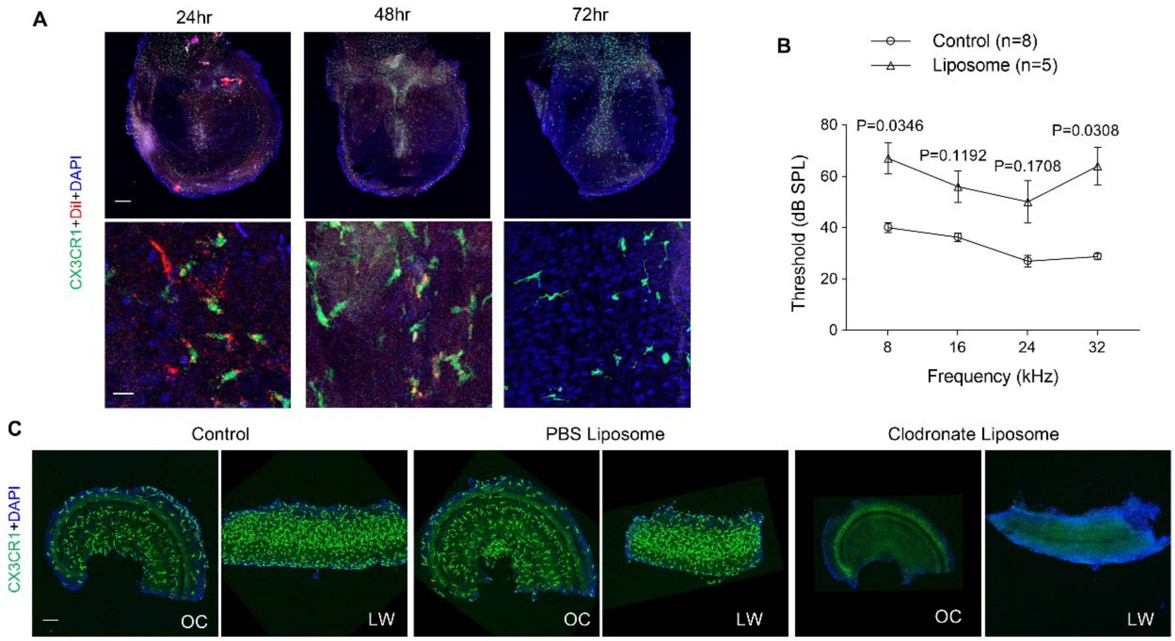


**Fig. S4. Clodronate Liposomes induced systemic macrophage-depletion in both eardrum and cochlea.** (**A**) Accumulation of Dil liposomes in eardrum macrophages (red arrows) at 24, 48 and 72 hours after facial intravenous injection. Scale bars, 200 µm (*top*), 20 µm (*bottom*). (**B**) ABR was measured at P21 and compared between normal (n=8) and macrophage depleted (n=5) groups at 8, 16, 24 and 32 kHz frequencies. (**C**) Representative confocal images of cochlear macrophages at P15 in normal, liposome control, and macrophage-depleted animals. Scale bar, 100 µm. (OC: Organ of Coti, LW, lateral wall). Data were presented as mean ± SEM, statistical significance was calculated through Two-way ANOVA with Šídák's test. ($F_{requency\ x\ Group}$ (3, 33) = 2.945, $F_{requency}$ (2.885, 31.73) = 10.01, $F_{Group}$ (1, 11) = 30.10, $F_{Threshold}$ (11, 33) = 6.071).

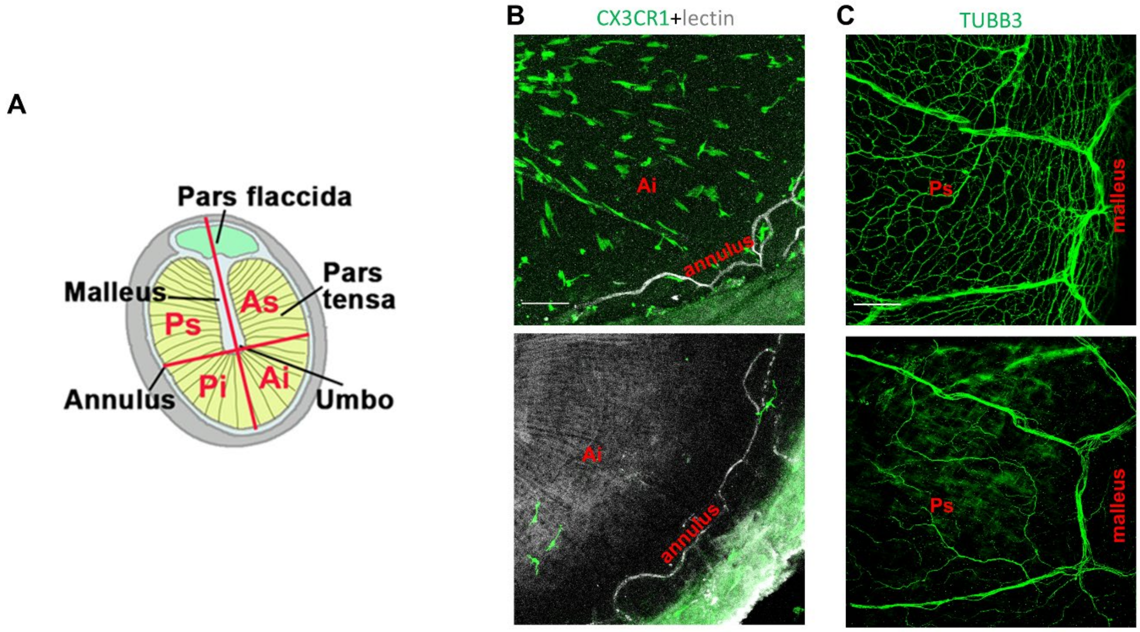


**Fig. S5. Depletion of Macrophages Leads to Underdeveloped Nerve Fibers**. (**A**) the illustration depicts the eardrum, which consists of two anatomical regions: the pars tensa and the pars flaccida. It is divided into four quadrants by an imaginary vertical line drawn along the manubrium of the malleus and another line that passes through the umbo, perpendicular to the first line. The quadrants are labeled as follows: anterosuperior (As), anteroinferior (Ai), posteroinferior (Pi), and posterosuperior (Ps). The thickness of the TM will be measured in these four areas. (**B, C**) The upper panels illustrate the distribution of macrophages and nerve fibers in the pars tensa near the annulus in control animals. The lower panels demonstrate that the nerve fibers in macrophage-depleted animals are underdeveloped, and some fibers are missing altogether. Scale bar: 100μm (*left*), 50μm (*right*).

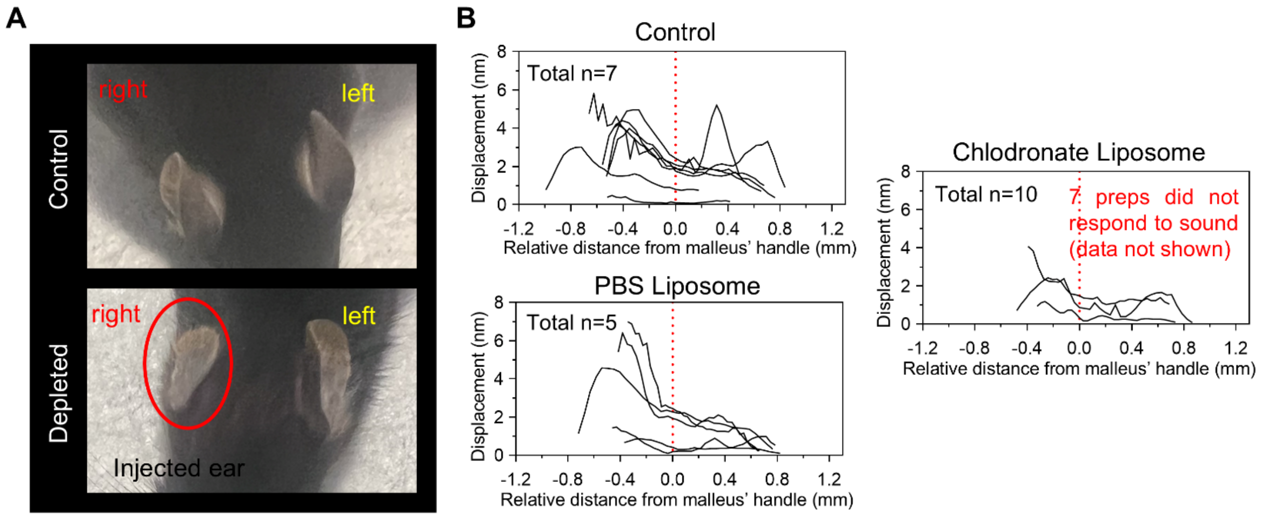


**Fig. S6. Depletion of Macrophages Leads to morphology and OCT function change**. (**A**) Local depletion of Macrophages Leads to smaller and "soft" pinna (**B**) Measurements of eardrum displacement (systemic cohort) as a function of horizontal distance from the malleus' handle in reaction to an 8 kHz 81 dB SPL sound stimulus.

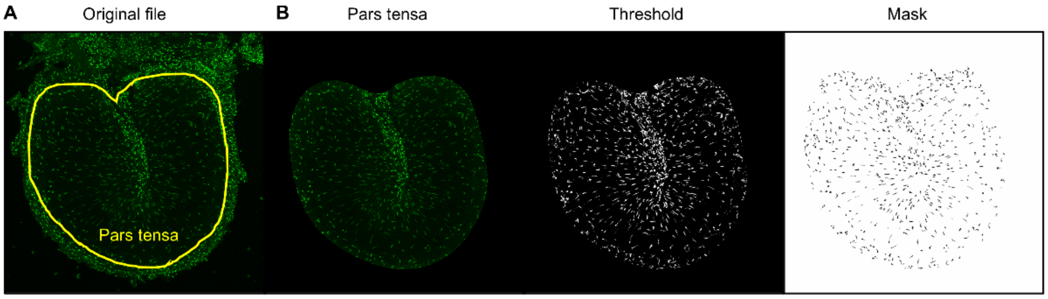


**Fig. S7**. **Macrophage quantification and measurements of the eardrum area were conducted using Fiji (ImageJ, version 1.54p).** (**A**) Displays the projected 2D representations of the tympanic membrane, created by stitching together and Z-projecting the image stacks. (**B**) Illustrates how each image was individually thresholded and masked using the "Analyze Particles" function in Fiji.
